# The LRRC8A:C Heteromeric Channel Is a cGAMP Transporter and the Dominant cGAMP Importer in Human Vasculature Cells

**DOI:** 10.1101/2020.02.13.948273

**Authors:** Lauren J. Lahey, Xianlan Wen, Rachel E. Mardjuki, Volker Böhnert, Gaelen T. Hess, Christopher Ritchie, Jacqueline A. Carozza, Merritt Maduke, Michael C. Bassik, Lingyin Li

## Abstract

Extracellular 2’3’-cyclic-GMP-AMP (cGAMP) is an immunotransmitter secreted by cancer cells and taken up by host cells to activate the anti-cancer STING pathway. No cGAMP exporter has been identified, and SLC19A1, a recently identified cGAMP importer, does not account for the import activity in most cell types. Here, we identify the LRRC8A:C heteromeric channel, a volume-regulated anion channel (VRAC), as a cGAMP transporter. This channel mediates cGAMP import or export depending on the cGAMP chemical gradient, and channel activation or inhibition modulates cGAMP transport. Other 2’3’-cyclic dinucleotides are also transported by LRRC8A:C channels, including the investigational cancer therapeutic ADU-S100. Furthermore, we demonstrate that the LRRC8A-containing channel is the dominant cGAMP importer in primary human vasculature cells. Given tumor vasculature’s regulation of immune infiltration and its disruption in response to STING agonists, we have uncovered a leading molecular mechanism for extracellular cGAMP signaling in this important anti-cancer target.

## INTRODUCTION

A potent and versatile immune signaling molecule, 2’3’-cyclic-GMP-AMP (cGAMP) is a second messenger that couples detection of pathogen- or damage-associated threats to activation of innate immunity. cGAMP is produced when mis-localized DNA is recognized in the cytosol by cyclic-AMP-GMP synthase (cGAS) in metazoan cells (Ablasser et al., 2013a; Diner et al., 2013; Gao et al., 2013; Wu et al., 2013; Zhang et al., 2013). Cytosolic double-stranded DNA (dsDNA) represents a conserved danger signal that can arise from diverse threats to a cell: dsDNA viruses (Li et al., 2013), retroviruses (Gao et al., 2015), bacteria (Manzanillo et al., 2012; Nandakumar et al., 2019), damage (Harding et al., 2017; Hatch et al., 2013; McArthur et al., 2018), cancerous states (Bakhoum et al., 2018; Mackenzie et al., 2017), and senescence (Dou et al., 2017; Glück et al., 2017; Yang et al., 2017). Once synthesized, cGAMP activates its receptor, Stimulator of Interferon Genes (STING) (Ablasser et al., 2013a; Diner et al., 2013; Zhang et al., 2013), to trigger activation of TBK1 and IRF3 signaling (Liu et al., 2015; Tanaka and Chen, 2012; Zhong et al., 2008). The resulting production of type I interferons and other cytokines then induces powerful inflammatory and defense responses (Sun et al., 2013). However, aberrant activation of this pathway, such as from DNAse deficiencies (Gao et al., 2015; Stetson et al., 2008) or mitochondrial damage (McArthur et al., 2018), can lead to deleterious effects such as autoimmune symptoms (Crowl et al., 2017). In order to understand the crucial roles of cGAS-STING signaling in homeostasis and disease, it is imperative to determine how nature regulates the potent central messenger of the system, cGAMP, including its trafficking and transport.

cGAMP is not only a cell-intrinsic activator of STING, but also a paracrine signal that orchestrates larger-scale biological responses. During viral infection, cGAMP produced in infected cells passes through connexin gap junctions into immediately adjacent cells to activate local anti-viral innate immunity (Ablasser et al., 2013b). The crucial importance of cGAMP transfer was recently highlighted even when exogenously administered as an intranasal influenza vaccine adjuvant. Although cGAMP-containing pulmonary surfactant biomimetic nanoparticles are first taken up by alveolar macrophages, subsequent intercellular transfer of cGAMP to alveolar epithelial cells and STING activation in these stromal cells are required for generation of heterosubtypic antiviral immunity (Wang et al., 2020). Recently, we reported a new cGAMP paracrine signaling mechanism: cGAMP is directly exported into the extracellular space in a soluble, non-membrane bound form (Carozza et al., 2020). We demonstrated that cancer cells basally produce and export cGAMP. Secreted extracellular cGAMP from cancer leads to an increase in innate immune cells within the tumor microenvironment and contributes to the curative effect of ionizing radiation in a STING-dependent manner (Carozza et al., 2020). While the importance of extracellular cGAMP signaling has thus far been demonstrated in the context of anti-cancer immunity (Zhou et al., 2020), it could, conceivably, operate in numerous physiological settings.

Despite this newly discovered role of cGAMP as an extracellular messenger, there remains a large gap in knowledge as to how the molecule is exported and imported by cells. cGAMP is negatively charged and must rely on facilitated mechanisms to cross the plasma membrane. We and others identified the reduced folate carrier SLC19A1 as the first importer of extracellular cGAMP (Luteijn et al., 2019; Ritchie et al., 2019a), however our studies clearly indicated the presence of additional cGAMP import mechanisms. We were unable to identify a primary human cell type that uses SLC19A1 as its dominant cGAMP importer (Ritchie et al., 2019b, 2019a). Our previous work revealed that different cell types have varying degrees of sensitivity to extracellular cGAMP and that part of this difference appears to be controlled through the use of different import machinery. Furthermore, the identity of the cGAMP exporter(s) in cancer cells is currently unknown. Discovery of additional cGAMP transport mechanisms is therefore imperative to understand the extent, selectivity, and regulation of extracellular cGAMP signaling in homeostasis and disease.

Here, through genetic, pharmacologic, and electrophysiological approaches, we identified the LRRC8A:C heteromeric channel as a second direct transporter of cGAMP. Interestingly, the transmembrane pore of LRRC8A channels shares structural similarities with that of connexins—known gap junction cGAMP transporters—however LRRC8A channels open to the extracellular space instead of linking together the cytosol between two cells (Deneka et al., 2018). LRRC8A was recently identified as an essential component of the long sought-after volume-regulated anion channel (VRAC) (Qiu et al., 2014; Voss et al., 2014), but its role appears to extend beyond regulation of cell-intrinsic physiology as LRRC8A-containing channels are now being discovered to mediate cell-cell communication via transport of signaling molecules (Chen et al., 2019). We now identify immunostimulatory 2’3’-cyclic dinucleotides as a new substrate class for LRRC8A channels and demonstrate cGAMP import or export depending on the chemical gradient and activation state of the channel. In establishing a connection between the burgeoning fields of cGAMP signaling in innate immunity and LRRC8A channel physiology, new avenues of research into the biological relationship between these processes and the development of therapeutics that target this cGAMP signaling mechanism are opened.

## RESULTS

### A Genome-Wide CRISPR Screen Identifies LRRC8A as a Positive Regulator of Extracellular cGAMP-Mediated STING Pathway Activation

We previously identified SLC19A1 as the first cGAMP importer by performing a whole-genome CRISPR knockout screen in the U937 monocyte-derived cell line (Ritchie et al, 2019). While SLC19A1 was the dominant importer in U937 cells, the existence of additional cGAMP import pathways was evident from the residual signaling observed in *SLC19A1*^*−/−*^ cells: extracellular cGAMP treatment still led to activation of STING and downstream phosphorylation of STING, the kinase TBK1, and the transcription factor IRF3, ultimately resulting in cytokine production and cell death (Figure 1A) (Ritchie et al., 2019a). To identify these SLC19A1-independent import mechanisms, we performed a whole-genome CRISPR knockout screen (Morgens et al., 2017) in U937 Cas9-*SLC19A1*^*−/−*^ cells. We determined that ~30 μM extracellular cGAMP was a 50% lethal dose (LD_50_) in U937 Cas9-*SLC19A1*^*−/−*^ cell at 48 hours (Figure S1C), suggesting that our previous live/dead screen approach could again be employed. We treated the library daily for 12 days with cGAMP, increasing from 15 μM to 30 μM, while passaging untreated cells in parallel as controls (Figure 1B). In principle, cells harboring an sgRNA that knocks out a positive regulator of the extracellular cGAMP-STING pathway, including a cGAMP importer, should be enriched for via resistance to cGAMP-mediated cell death. Conversely, cells should be depleted if they harbor sgRNAs targeting negative regulators of the pathway. At the end of the selection, we isolated genomic DNA from each condition and sequenced the encoded sgRNAs. We then analyzed fold changes of sgRNA sequences in cGAMP treated cells versus controls using the Cas9 high-Throughput maximum Likelihood Estimator (casTLE) statistical framework to calculate a casTLE score and effect for each gene (Morgens et al., 2016). Key STING pathway components, *TMEM173* (STING), *TBK1*, and *IRF3*, were reproducibly identified with high confidence scores (Figure 1C) and large effect sizes (Figure 1D), validating this screening approach. Interestingly, *LRRC8A* clustered near known STING pathway members (Figure 1C – 1D).

**Figure 1.**
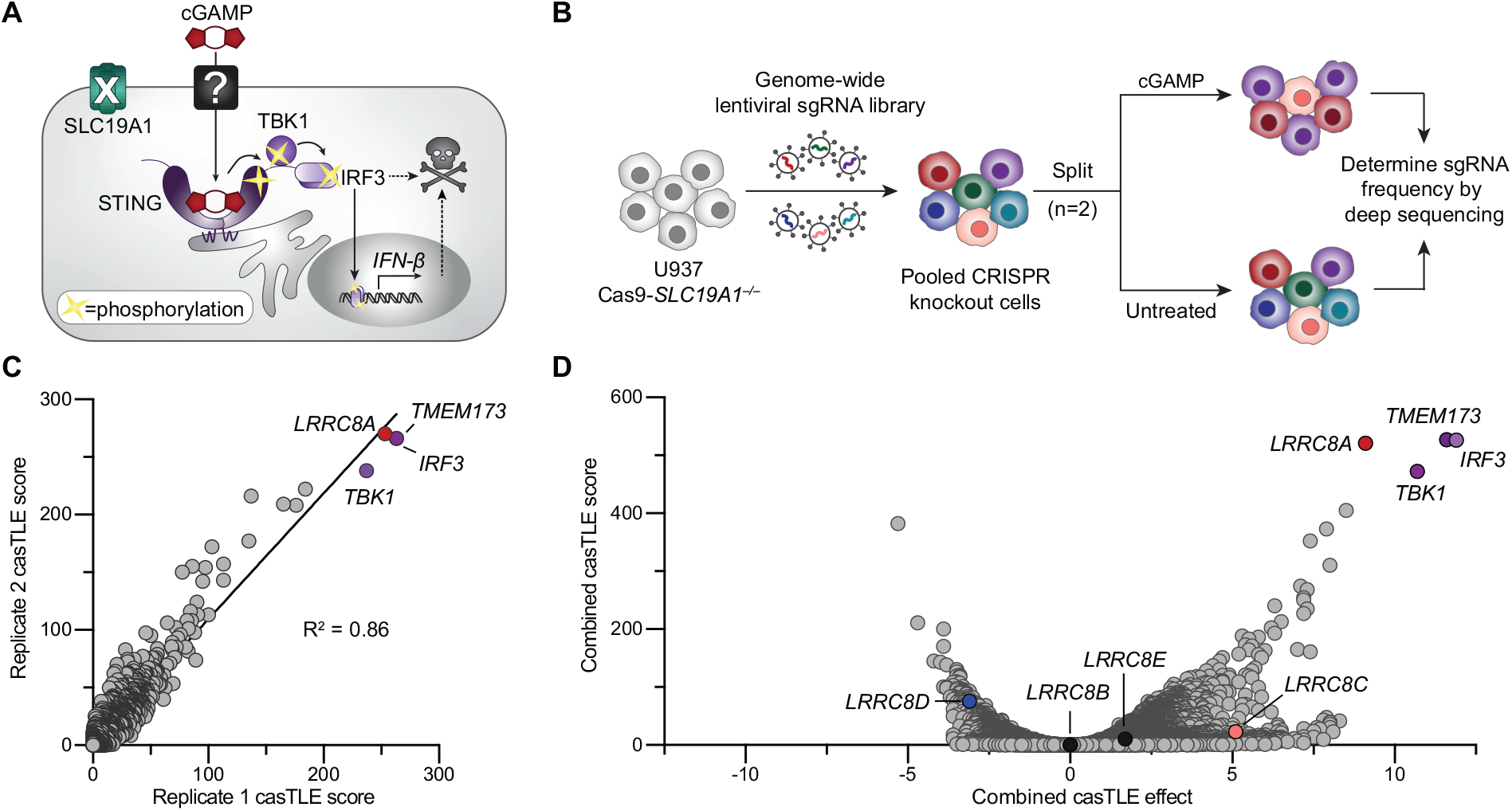
A Genome-Wide CRISPR Screen Identifies LRRC8A as a Positive Regulator of Extracellular cGAMP-Mediated STING Pathway Activation. (A) Scheme of extracellular cGAMP-STING signaling in U937 *SLC19A1*^−/−^ cells. cGAMP imported from the extracellular space binds to and activates STING. The resulting signaling cascade leads to phosphorylation of STING, the kinase TBK1, and the transcription factor IRF3, ultimately leading to cytokine production and cell death. (B) CRISPR screen strategy. U937 Cas9-*SLC19A1*^−/−^ cells were transduced with a whole-genome sgRNA lentiviral library. In two replicates, library cells were treated with cGAMP or untreated for 12 days. Genomic DNA was harvested, deep sequenced, and analyzed for sgRNA depletion or enrichment. (C) Plot of casTLE score for every gene in replicate 1 versus replicate 2 with displayed correlation across all genes (R^2^ = 0.86). Top scoring hits are annotated. (D) Volcano plot of casTLE effect size versus score for all genes across combined replicates. Top enriched hits, as well the *LRRC8* family, are annotated. See also Figure S1.

### LRRC8A and LRRC8 Paralogs Differentially Facilitate cGAMP Import

LRRC8A was recently identified as an essential component of the volume-regulated anion channel (VRAC) that resides on the plasma membrane (Qiu et al., 2014; Voss et al., 2014). Upon cell swelling or sensing of various stimuli, opening of VRAC leads to outward flow of chloride ions, organic osmolytes, and water to facilitate regulated volume decrease (Chen et al., 2019; Osei-Owusu et al., 2018; Strange et al., 2019). In order to form channels with functional VRAC activity in cells, LRRC8A must associate with one or more paralogous proteins, LRRC8B–E (Voss et al., 2014). These channels are reported to be heteromeric hexamers in which subunit stoichiometry may be variable (Gaitán-Peñas et al., 2016; Lutter et al., 2017). Depending on the paralog(s) in complex with LRRC8A, the channel exhibits different properties (Syeda et al., 2016) and can transport different substrates. While complexes containing LRRC8B–E mediate inorganic anion flux, LRRC8A:C, LRRC8A:D, and LRRC8A:E complexes transport larger anionic substrates such as glutamate and aspartate (Gaitán-Peñas et al., 2016; Lutter et al., 2017; Planells-Cases et al., 2015; Schober et al., 2017). LRRC8A:C and LRRC8A:E channels have been reported to transport ATP (Gaitán-Peñas et al., 2016). Distinct characteristics of LRRC8D-containing channels have been reported, including mediating efflux of uncharged cellular osmolytes (Schober et al., 2017) and import of the drugs blasticidin, cisplatin, and carboplatin (Lee et al., 2014; Planells-Cases et al., 2015). It is therefore plausible that LRRC8A forms heteromeric channels with other LRRC8 paralogs to import extracellular cGAMP.

In our screen, *LRRC8C* and *LRRC8E* had moderate and weak positive effect sizes, respectively, and *LRRC8D* had a negative effect size (Figure 1C). To investigate the role of all LRRC8 paralogs in extracellular cGAMP signaling, we individually knocked out each gene in U937 *SLC19A1*^*−/−*^ cells using multiple gene-specific sgRNAs. We also used a non-targeting scrambled sgRNA to serve as a negative control. We then isolated and validated single cell clones, treated them with extracellular cGAMP, and measured phosphorylation of IRF3 (p-IRF3) to assess STING pathway activation (Figure S2).

LRRC8A knockout clones exhibited a ~40% reduction in p-IRF3 relative to total IRF3 as compared to scramble controls, while LRRC8C knockout clones exhibited a ~30% reduction (Figure 2A). In contrast, a ~40% increase in p-IRF3 signaling was observed in LRRC8D knockout clones. Negligible differences were observed in LRRC8B and LRRC8E knockouts (average decreases of 5% and 8%, respectively), consistent with LRRC8B not playing a role in transport of large anions (Lutter et al., 2017; Schober et al., 2017) and LRRC8E expression being restricted (Chen et al., 2019). Together, these data validate the effects observed in the whole-genome screen that LRRC8 paralogs play differential regulatory roles in extracellular cGAMP signaling.

**Figure 2.**
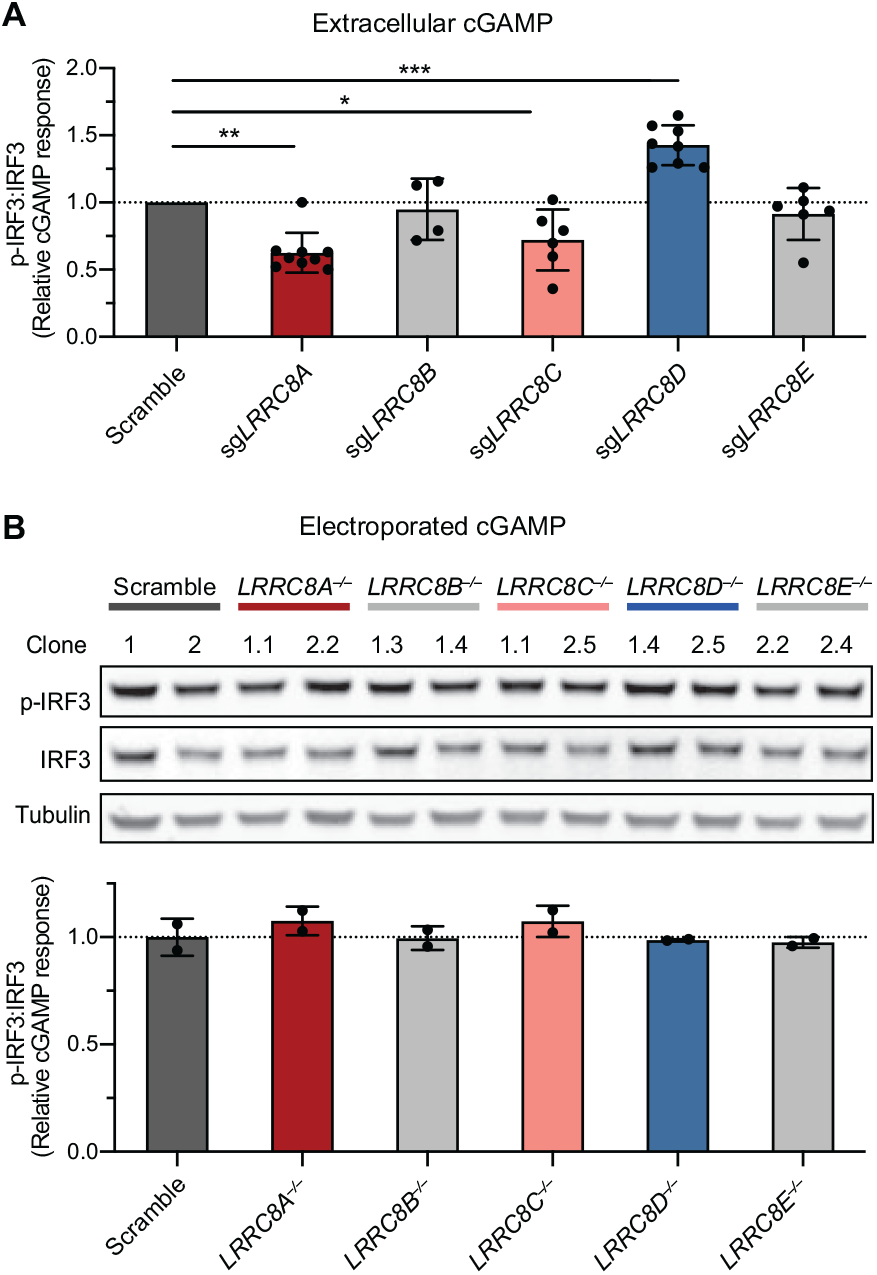
LRRC8A and LRRC8 Paralogs Differentially Facilitate cGAMP Import. (A) Effect of LRRC8A-E on extracellular cGAMP signaling. U937 *SLC19A1*^*−/−*^ clones expressing scramble or *LRRC8A*-*LRRC8E* sgRNAs were treated with 100 μM cGAMP for 4 h and signaling was assessed by Western blot (see Figure S2). Summarized results for clones (confirmed heterozygous or homozygous knockout) are plotted relative to the respective scramble control average (n = 4-9 biological replicates). Significance calculated using two-tailed *t*-test; \**P* ≤ 0.05, \*\**P* ≤ 0.01, \*\*\**P* ≤ 0.001. (B) Effect of LRRC8A-LRRC8E on intracellular cGAMP signaling. U937 *SLC19A1^−/−^-*scramble or −*LRRC8A*^−/−^ through −*LRRC8E*^−/−^ clones were electroporated with 100 nM cGAMP, cultured for 2 h, and assessed for signaling by Western blot (n = 2 biological replicates). For (A)-(B), results plotted as mean ± SD. See also Figure S2.

We then tested whether LRRC8 paralogs affect the response to cGAMP at the level of import or by altering downstream STING pathway signaling. When cGAMP was electroporated into the cells, bypassing the need for specific importers, *LRRC8A*^*−/−*^ through *LRRC8E*^*−/−*^ clones no longer exhibited differential p-IRF3 signaling compared to scramble controls (Figure 2B). Our data support a model in which LRRC8A, likely in complex with LRRC8C, facilitates transport of extracellular cGAMP across the plasma membrane. Our data also demonstrate that LRRC8D negatively affects this process, possibly by sequestering LRRC8A into heteromeric complexes that are not functional for mediating cGAMP import.

### LRRC8A-Containing VRAC Channels Directly Transport cGAMP

We next sought to determine whether cGAMP is directly transported by LRRC8A channels. Alternatively, VRAC activation could alter membrane potential to influence cGAMP transport through another route not revealed by our CRISPR screen. Much of the work dissecting VRAC function has been performed in HEK293 cell lineages (Osei-Owusu et al., 2018; Pedersen et al., 2015), which we previously determined respond to extracellular cGAMP in an SLC19A1-independent manner (Ritchie et al., 2019a). Following the generation of LRRC8 knockout pools and validation of gene editing (Figure S3A), we tested whether these channels facilitate cGAMP import in HEK293 cells. Using the most upstream reporter of cGAMP import, phosphorylation of STING (p-STING) following cGAMP binding, we observed that HEK293 LRRC8A knockout pools exhibited a ~20% decrease in cellular response to extracellular cGAMP compared to controls (Figure 3A). Conversely, LRRC8D knockout pools exhibited a ~30% increase in extracellular cGAMP response. These results suggest that LRRC8 channels also facilitate import of cGAMP in HEK293 cells, although they do not account for the dominant mechanism. Consistent with our observations in U937 cells, LRRC8 paralogs differentially affect the cGAMP import process in HEK293 cells.

**Figure 3.**
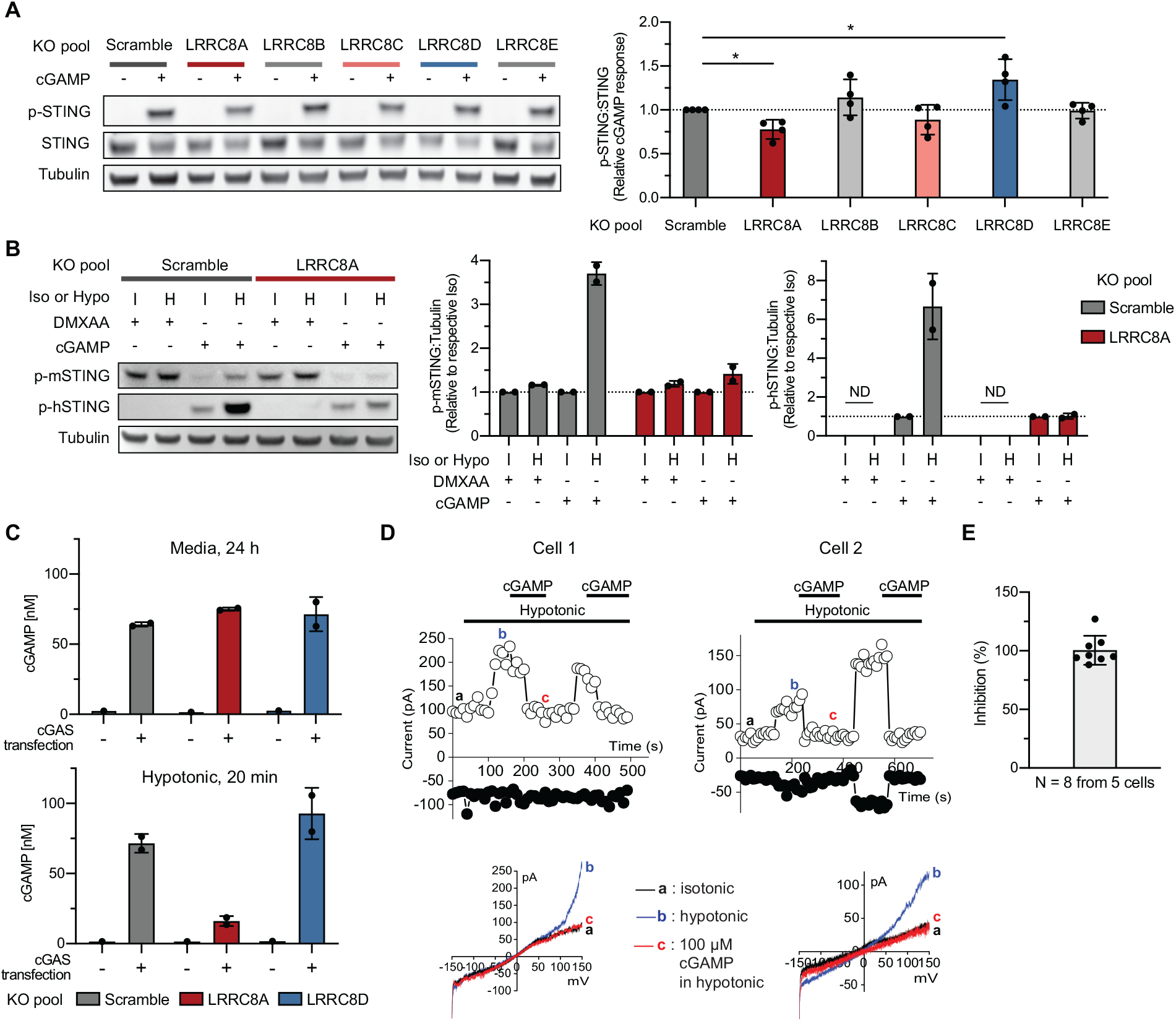
LRRC8A-Containing VRAC Channels Directly Transport cGAMP. (A) Effect of LRRC8A-E on extracellular cGAMP signaling in HEK293 cells. HEK293 scramble or LRRC8A-LRRC8E knockout pools were treated with 100 μM cGAMP for 2 h and signaling assessed by Western blot. Representative blots displayed with quantification (n = 4 biological replicates). (B) Effect of osmolarity on DMXAA and cGAMP signaling in HEK293 cells. Scramble or LRRC8A knockout cells were transfected for expression of mouse STING. After 24 h, cells were treated with 250 μM DMXAA or 100 μM GAMP in isotonic or hypotonic buffer for 1 h. Activation of mouse (p-mSTING) or human (p-hSTING) signaling was assessed by Western blot. Representative blots displayed with quantification (n = 2 technical replicates). ND = not detected. (C) Extracellular cGAMP concentrations from HEK293 scramble, LRRC8A, or LRRC8D knockout pools were measured by LC-MS/MS following cGAS plasmid transfection from media (24 h) or after hypotonic buffer stimulus (20 min) (n = 2 technical replicates). (D) Representative whole-cell patch clamp current traces measured in HEK293 cells upon hypotonic stimulation in the absence or presence of cGAMP (100 μM). Overlays of the current-voltage relationship at the indicated points are also displayed. For Additional traces from three separate cells, Figure S3. (E) Summary data for inhibition of VRAC current by 100 μM extracellular cGAMP. Inhibition of hypotonic-induced current at +150 mV is displayed (n = 8 measured in 5 cells). For (A), (B), (C), and (E), data are mean ± SD. Significance calculated using ratio paired *t*-test; \**P* ≤ 0.05. See also Figure S3.

We hypothesized that if cGAMP is a substrate of LRRC8A, further opening of the channel would increase import of extracellular cGAMP and downstream STING signaling. Activation of VRAC is commonly experimentally achieved by application of hypotonic extracellular buffer (Qiu et al., 2014; Voss et al., 2014). Furthermore, we predicted that the membrane-permeable STING agonist, DMXAA, would act independently of LRRC8A opening. It is important to note that DMXAA activates murine, but not human, STING, while cGAMP activates both (Conlon et al., 2013; Kim et al., 2013). HEK293 cells, which express low levels of human STING, were therefore transfected for co-expression of mouse STING. Cells were treated with either isotonic (control) or hypotonic (activating) solutions containing either DMXAA or cGAMP. DMXAA treatment in hypotonic buffer led to modest increases in mouse p-STING relative to isotonic treatment in both scramble and LRRC8A knockout HEK293 cells (16% and 19%, respectively) (Figure 3B). In contrast, cGAMP treatment in hypotonic buffer resulted in a ~250% increase in mouse p-STING relative to isotonic treatment in scramble control cells, and this increase was mostly abolished in LRRC8A knockout pools. As expected, no activation of human STING was observed with DMXAA treatment. LRRC8A-dependent hypotonic potentiation of cGAMP signaling was also detected through measurement of human p-STING, with a nearly 600% increase observed in scramble control cells but only an 11% increase in the LRRC8A knockout pool. These results demonstrate that LRRC8A channel activation boosts signaling for one STING agonist (cGAMP) but not another (DMXAA), in line with their differential need for facilitated membrane transport.

If LRRC8A channels directly transport cGAMP, we should be able to observe cGAMP efflux by i) inverting the cGAMP gradient to a high intracellular concentration and ii) triggering VRAC opening. It is worth noting that VRAC-mediated export of another nucleotide, ATP, has been demonstrated upon channel opening (Dunn et al., 2020; Gaitán-Peñas et al., 2016; Hisadome et al., 2002). We stimulated intracellular cGAMP synthesis in HEK293 scramble control, LRRC8A, or LRRC8D knockout pools by transfecting cells with a cGAS expression plasmid. Degradation of cGAMP by its hydrolase, ENPP1, was inhibited using the small molecule inhibitor STF-1084 (Carozza et al., 2020). After 24 hours, we harvested the media from each condition and treated cells with a hypotonic buffer for 20 minutes to stimulate channel opening. We then measured cGAMP concentrations in the extracellular solutions. We detected cGAMP secreted into the media at levels similar to those previously reported, but observed no notable differences between scramble, LRRC8A, and LRRC8D knockout pools (64, 75, and 71.4 nM, respectively), suggesting that LRRC8A channels are not major exporters of cGAMP in HEK293 cells under resting conditions (Figure 3C). In contrast, we detected markedly higher cGAMP export from control (72 nM) and LRRC8D knockout cells (93 nM) compared to LRRC8A knockout cells (16 nM) upon hypotonic treatment (Figure 3C). These data suggest that LRRC8A-containing channels, when opened, mediate cGAMP export.

We next tested whether cGAMP directly interacts with LRRC8A channels. Previous work using electrophysiology has shown that extracellular ATP inhibits Cl^-^ influx via VRAC and that this inhibition is the result of direct pore block (Jackson and Strange, 1995; Kefauver et al., 2018; Tsumura et al., 1996). We tested whether extracellular cGAMP also blocks Cl^-^ influx through VRAC. Whole-cell patch clamp analysis of HEK293 cells revealed that the characteristic, hypotonic solution-induced VRAC current measured at positive membrane potential is inhibited by introduction of cGAMP in the bath solution (Figure 3D, 3E, and Figure S3B). Similar to inhibition by ATP (Figure S3C), inhibition by cGAMP is rapid and reversible, providing strong support for the conclusion that cGAMP directly interacts with the LRRC8A-containing VRAC channel.

The observed inhibition is also consistent with cGAMP being directly transported by the LRRC8A-containing channels: larger substrates that are transported slowly generally act as inhibitors (“permeating blockers”) of faster-traveling, small substrates through channels (Woodhull, 1973) – ATP was previously shown to be a such a permeating blocker of Cl^-^ influx through VRAC (Hisadome et al., 2002). Therefore, it is likely that cGAMP, like ATP, permeates through the LRRC8A channel.

### The LRRC8A:C Channel Is the Dominant cGAMP Importer in Microvasculature Cells

The LRRC8 subunit analysis performed in U937 and HEK293 cells predicts that cell types expressing high levels of LRRC8A and LRRC8C and low levels of LRRC8D should import cGAMP effectively. We evaluated RNA expression profiles of different cell types in The Human Protein Atlas (proteinatlas.org) and identified that several vasculature cell lineages fit such a profile (Figure S4A). TIME cells, an immortalized human microvascular endothelial cell (HMVEC) line, express *LRRC8A* and *LRRC8C* transcripts at 15.2-fold and 1.6-fold higher levels, respectively, than U937 cells while expressing *LRRC8D* transcripts at a 4.3-fold lower level. Indeed, TIME cells responded to extracellular cGAMP with an EC_50_ of ~24 μM (Figure S4B), which is more sensitive than most cell lines we have evaluated to date, including U937 cells with an EC_50_ of 270 μM (Ritchie et al., 2019a). To determine if cGAMP import depends on LRRC8 complexes in these cells, we knocked out LRRC8A–E using CRISPR, as well as the first identified importer, SLC19A1 (Figure S4C). LRRC8A or LRRC8C knockout in TIME cells resulted in loss of the majority of response to extracellular cGAMP, as measured by phosphorylation of all key pathway components: p-STING, p-TBK1, and p-IRF3 (Figure 4A). LRRC8D knockout cells trended towards a modestly increased p-STING response, the amplitude of which is lower than in U937 or HEK293 cells, likely reflective of lower *LRRC8D* expression in TIME cells (Figure S4A). No significant effect was observed upon SLC19A1 knockout, confirming that this importer is not broadly used and that cGAMP import mechanisms are cell-type specific. Bypassing the need for specific importers using lipid-based transfection of cGAMP into cells resulted in similar responses across scramble, LRRC8A, LRRC8C and LRRC8D knockout pools (Figure 4B), again supporting a role for LRRC8A and LRRC8C in transport of cGAMP across the plasma membrane. Knockout of both LRRC8A and LRRC8C did not have an additive effect compared to single knockouts (Figure 4C). Given the known requirement of LRRC8A and at least one other subunit for cellular VRAC function (Syeda et al., 2016; Voss et al., 2014), our results are consistent with LRRC8A and LRRC8C forming heterometric complexes to import cGAMP.

**Figure 4.**
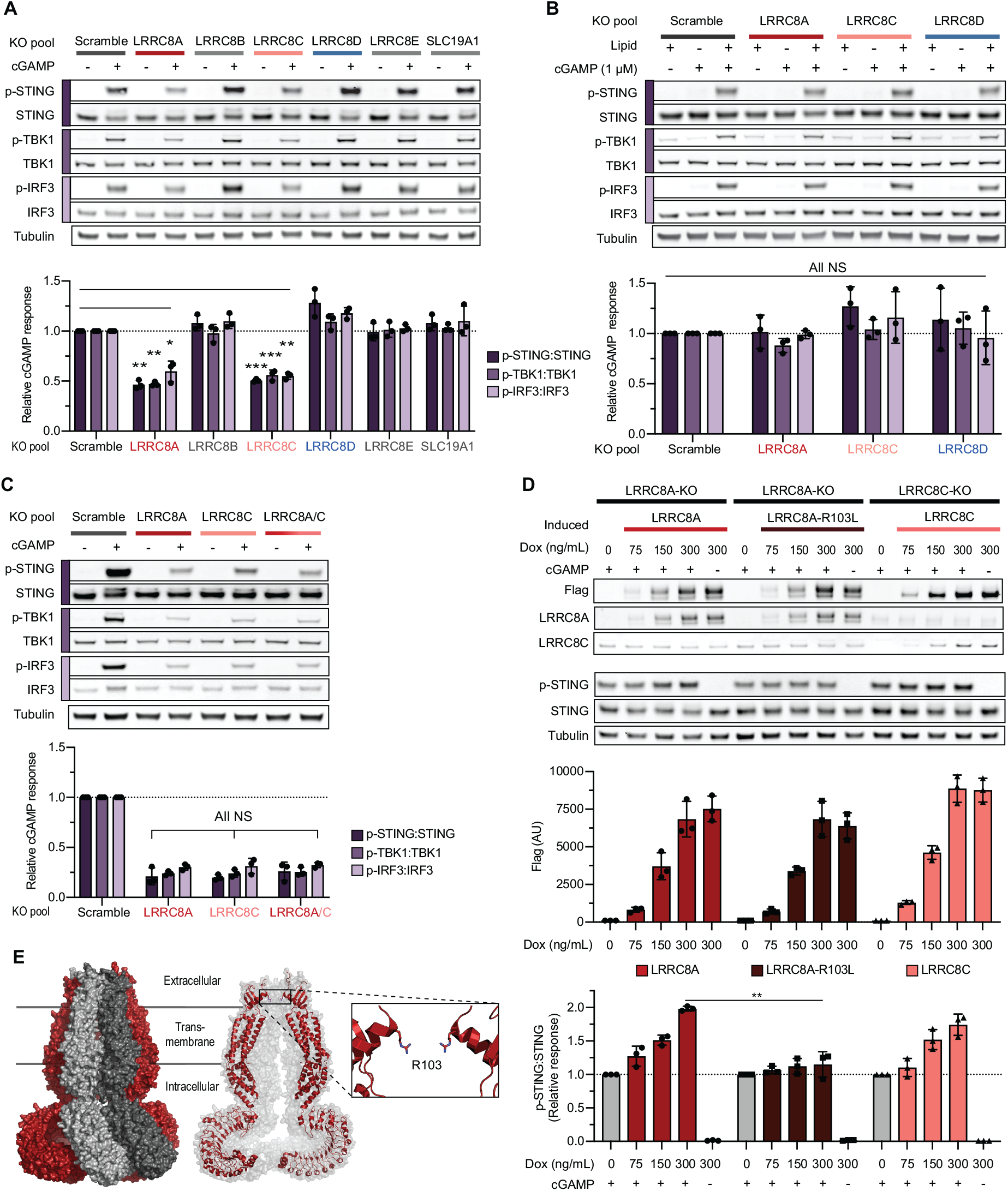
The LRRC8A:C Channel Is the Dominant cGAMP Importer in Microvasculature Cells. (A) Effect of LRRC8A-E and SLC19A1 on extracellular cGAMP signaling in TIME cells. Knockout pools were treated with 50 μM cGAMP for 2 h and signaling then assessed by Western blot (n = 3 biological replicates). (B) Effect of LRRC8A, LRRC8C, and LRRC8D on intracellular cGAMP signaling in TIME cells. Knockout pools were treated with 1 μM free or lipid-complexed cGAMP for 2 h and signaling was assessed by Western blot (n = 3 biological replicates). (C) Effect of LRRC8A, LRRC8C, and LRRC8A/C on extracellular cGAMP signaling in TIME cells. Knockout pools were treated with 50 μM cGAMP for 2 h and signaling then assessed by Western blot (n = 3 biological replicates). (D) Effect of LRRC8A, LRRC8A-R103L, or LRRC8C protein rescue on extracellular cGAMP signaling in TIME cells. Knockout cells were induced to express Flag-tagged protein constructs using doxycycline, then treated with 50 μM cGAMP for 2 h and signaling assessed by Western blot (n = 3 biological replicates). (E) Cryo-EM structure of human homo-hexameric LRRC8A (PDB: 5ZSU) (Kasuya et al., 2018). Surface representation of the complex, with cartoon representation of two opposing LRRC8A subunits and zoom to the extracellular pore gating R103 residue. For (A)-(D), representative blots are shown with quantification of all experiments displayed as mean ± SD. Significance calculated using ratio paired *t*-test; \**P* ≤ 0.05, \*\**P* ≤ 0.01, \*\*\**P* ≤ 0.001, not significant (NS). See also Figure S4.

To demonstrate that the effect of the CRISPR knockout is due to loss of LRRC8A and LRRC8C proteins, but not off-target effects of the CRISPR procedures, we next tested whether rescued expression of each protein restored signaling in the corresponding knockout TIME cells. Standard expression by transient transfection would result in plasmid DNA activating cGAS, production of cGAMP, and high basal STING pathway activation. Therefore, we generated stable cell lines in which LRRC8A-Flag and LRRC8C-Flag expression is driven from a tetracycline-inducible promoter. Unfortunately, we were not able to overexpress LRRC8D to probe its inhibitory role in these cells as its expression was consistently low or undetectable. Regardless, we observed concentration-dependent increases in extracellular cGAMP response when we restored LRRC8A and LRRC8C protein expression (Figure 4D). As revealed by recent cryo-electron microscopy structures of LRRC8A homomeric complexes (Deneka et al., 2018; Kasuya et al., 2018; Kefauver et al., 2018), the narrowest region of the LRRC8A channel is gated by an arginine residue (R103), creating a strong positive charge potential at the extracellular pore (Figure 4E). LRRC8C has a leucine residue at the corresponding position. The phosphodiester ring of cGAMP is double-negatively charged, therefore we predicted that R103 contributed by the LRRC8A subunit is crucial for cGAMP recognition. To test this hypothesis, we overexpressed the LRRC8A-R103L mutant in LRRC8A knockout cells and observed that it did not rescue cellular response to extracellular cGAMP (Figure 4D). This result supports a role for the pore-gating residue R103 in cGAMP recognition in a manner that is consistent with charge-charge interactions regulating channel access. Collectively, these data demonstrate that the LRRC8A:C complex functions as the dominant cGAMP importer in TIME cells.

### cGAMP Import by the LRRC8A:C Channel Can Be Potentiated by Sphingosine-1-Phosphate and Inhibited by DCPIB

After establishing the dominance of the LRRC8A:C cGAMP import pathway observed in TIME cells, we next wanted to identify chemical tools to regulate its function. VRAC has been shown to be activated downstream of sphingosine-1-phosphate (S1P) signaling to one of its G protein-coupled receptors, S1PR1 (Burow et al., 2014). In addition, endothelial cells are exposed to high S1P levels in the blood under physiological conditions (estimated 1 μM total), and also produce S1P themselves (Cartier and Hla, 2019). TIME cells are cultured with 5% fetal bovine serum, which we estimate to contain a maximum of 50 nM S1P. We hypothesized that increasing S1P closer to physiological levels would further increase cGAMP uptake by TIME cells *in vitro*. Additionally, we sought to determine the effect of DCPIB (Figure 5A), a known small molecule VRAC inhibitor (Decher et al., 2001; Qiu et al., 2014), on the response to extracellular cGAMP. The recently solved structure of a LRRC8A homomeric complex with DCPIB revealed that the inhibitor binds within the narrow pore formed by R103, effectively acting like a “cork in the bottle” (Kern et al., 2019). Since cGAMP uptake also required R103, we anticipated that DCPIB should inhibit cGAMP import. Indeed, S1P increased extracellular cGAMP signaling by 2- to 3-fold in a dose- and LRRC8A-dependent manner (Figure 5B), but did not lead to STING signaling in the absence of cGAMP (Figure S5). Conversely, DCPIB significantly inhibited all LRRC8A-dependent extracellular cGAMP signaling under both basal and S1P-stimulated conditions. Together, our results suggest that LRRC8A:C channel uptake of cGAMP into TIME cells is potentiated by physiological concentrations of S1P. In addition, DCPIB can be used as a tool to block LRRC8A-dependent cGAMP import.

**Figure 5.**
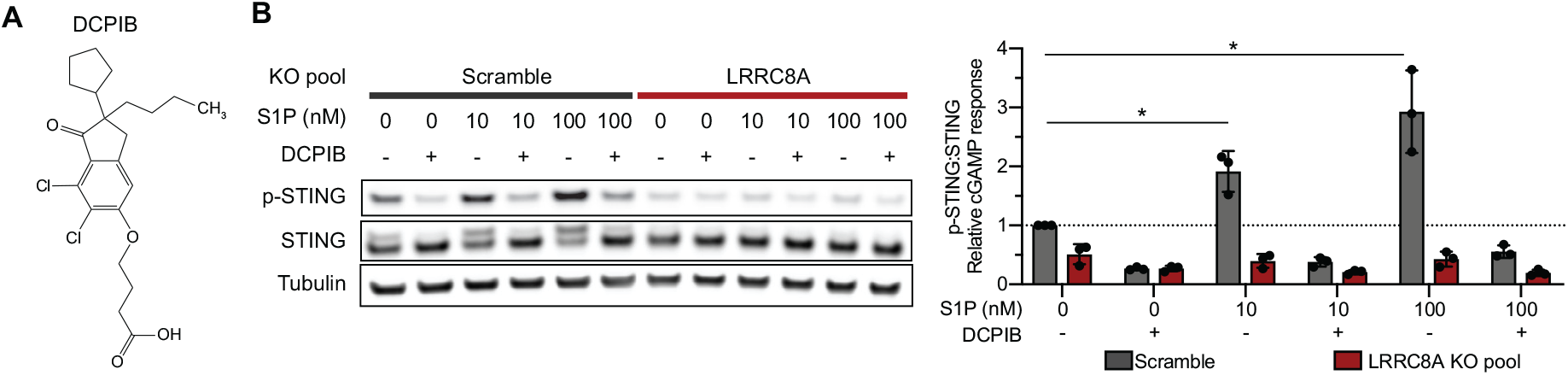
cGAMP Import by the LRRC8A:C Channel Can Be Potentiated by Sphingosine-1-Phosphate and Inhibited by DCPIB. (A) Chemical structure of DCPIB. (B) Effect of sphingosine-1-phosphate (S1P) and DCPIB on extracellular cGAMP signaling in TIME cells. Scramble and LRRC8A knockout pools were treated with cGAMP (50 μM), with or without S1P (10 or 100 nM) and DCPIB (20 μM) for 1 h and signaling assessed by Western blot (n = 3 biological replicates). Representative blots are shown with quantification of all experiments displayed as mean ± SD. Significance calculated using ratio paired *t*-test; \**P* ≤ 0.05. See also Figure S5.

### The LRRC8A:C Channel Imports Other 2’3’-Cyclic Dinucleotides

We next evaluated the specificity of LRRC8A complexes for import against a broad panel of natural and synthetic cyclic dinucleotides (CDNs) (Figure 6A). Synthetic non-hydrolyzable phosphorothioate cGAMP analogs 2’3’-cG^S^A^S^MP (Li et al., 2014) and 2’3’-CDA^S^ (Corrales et al., 2015) have been designed as potent STING agonists for innate immunotherapy. 2’3’-CDA^S^, also known as ADU-S100, is undergoing evaluation in clinical trials for treatment of metastatic cancers (ClinicalTrials.gov: NCT02675439, NCT03172936, and NCT03937141). Additionally, bacteria produce CDN second messengers with a different 3’3’-phosphodiester linkage between nucleotides. Bacterial CDNs include 3’3’-cyclic-GMP-AMP (3’3’-cGAMP), 3’3’-cyclic-di-GMP (3’3’-CDG), and 3’3’-cyclic-di-AMP (3’3’-CDA) (Burdette et al., 2011; Woodward et al., 2010; Zhang et al., 2013), which can trigger host STING signaling upon infection (Dey et al., 2015; Sauer et al., 2010; Woodward et al., 2010). We observed that LRRC8A channels account for the majority of cGAMP, 2’3’-cG^S^A^S^MP, and 2’3’-CDA^S^ import in TIME cells, while playing a minor role in 2’3’-CDA import and negligible role in the import of all 3’3’-CDNs (Figure 6B).

**Figure 6.**
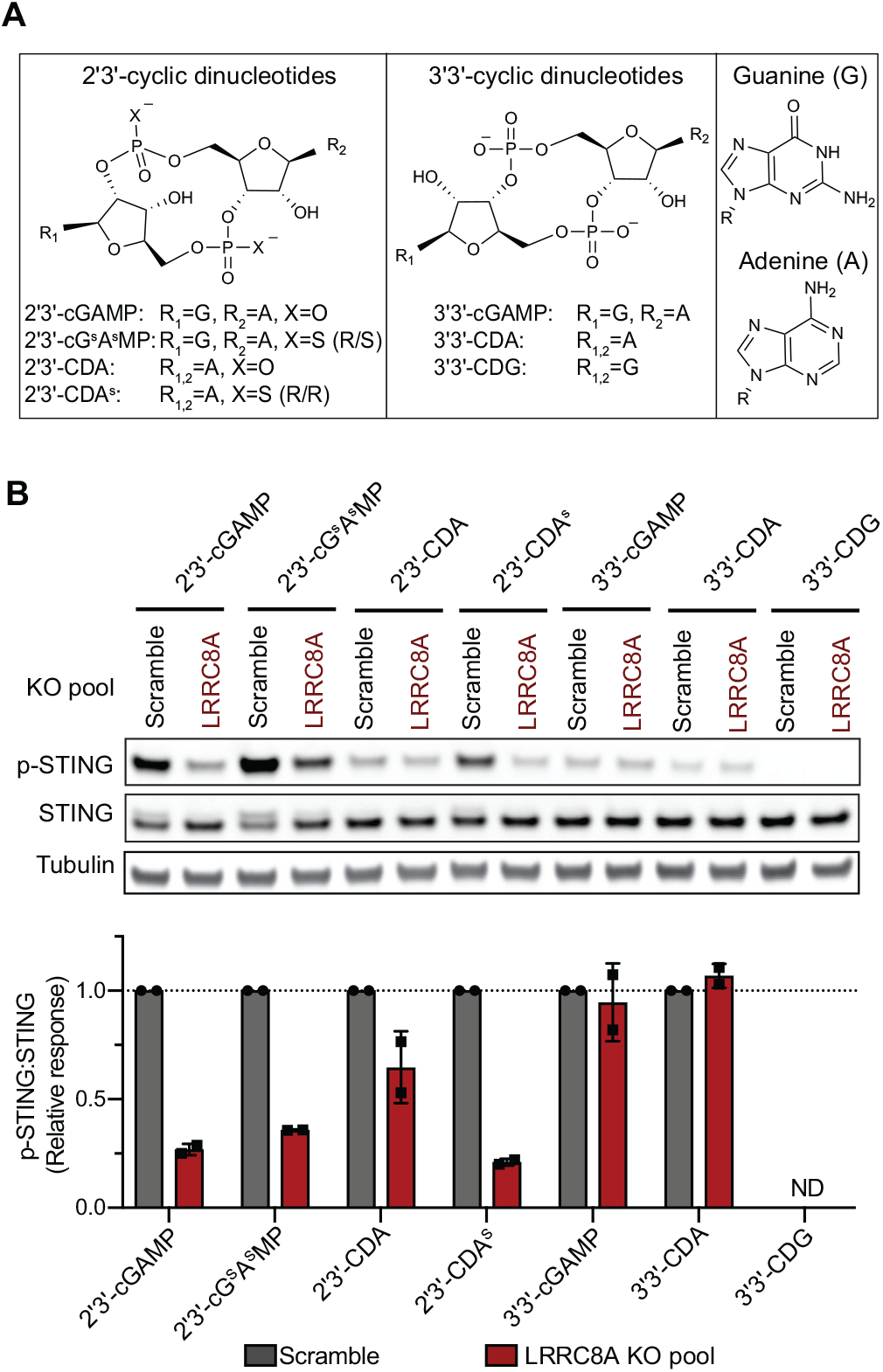
The LRRC8A:C Channel Imports Other 2’3’-Cyclic Dinucleotides. (A) Chemical structures of 2’3’- and 3’3’-cyclic dinucleotide (CDN) STING agonists. (B) Importance of LRRC8A for signaling of various extracellular CDNs in TIME cells. Scramble and LRRC8A knockout pools were treated with 50 μM cGAMP, 25 μM 2’3’-cG^S^A^S^MP, 100 μM 2’3’-CDA, 25 μM 2’3’-CDA^S^, 100 μM 3’3’-cGAMP, 200 μM 3’3’-CDA, or 200 μM 3’3’-CDG for 2 h and signaling assessed by Western blot (n = 2 biological replicates). Representative blots are shown with quantification of all experiments displayed as mean ± SD. ND = not detected.

### LRRC8A-Containing Channels Are the Dominant cGAMP Importer in Primary Human Endothelial Cells

Numerous studies have revealed that stromal (non-immune) cells are key responders to paracrine or pharmacologic STING activation and contribute to downstream anti-tumor signaling cascades. During early tumor engraftment in a B16.F10 melanoma model, spontaneous STING activation and robust interferon production are detected in endothelial cells prior to infiltration of dendritic cells and CD8^+^ T cells to the tumor microenvironment (Demaria et al., 2015). Through knockout and bone marrow chimera studies, it was found that STING signaling in the stroma, but not in the hematopoietic compartment, was required for tumor necrosis following injection of 2’3’-CDA^S^ in B16.F10 tumors (Francica et al., 2018). Therefore, elucidating the uptake mechanisms of extracellular cGAMP and its analogs into stromal lineages is necessary to understand the selectivity and regulation of this biologically important STING signaling axis.

The dominant import role of LRRC8A:C observed in TIME cells, an immortalized microvasculature endothelial lineage, strongly suggested that cGAMP import in primary human endothelial cells could also occur through LRRC8A channels. siRNA knockdown in primary human umbilical vein endothelial cells (HUVEC) decreased LRRC8A protein levels by ~70-90% (Figure 7A). As predicted, all siRNA treatments decreased extracellular cGAMP signaling, with oligo-9, −10, and −11 knocking down p-IRF3 signaling by more than 50%. These results demonstrate that the LRRC8A channel is not only the dominant cGAMP importer in immortalized human microvasculature cells, but also in primary HUVECs, and possibly in tumor vasculature.

**Figure 7.**
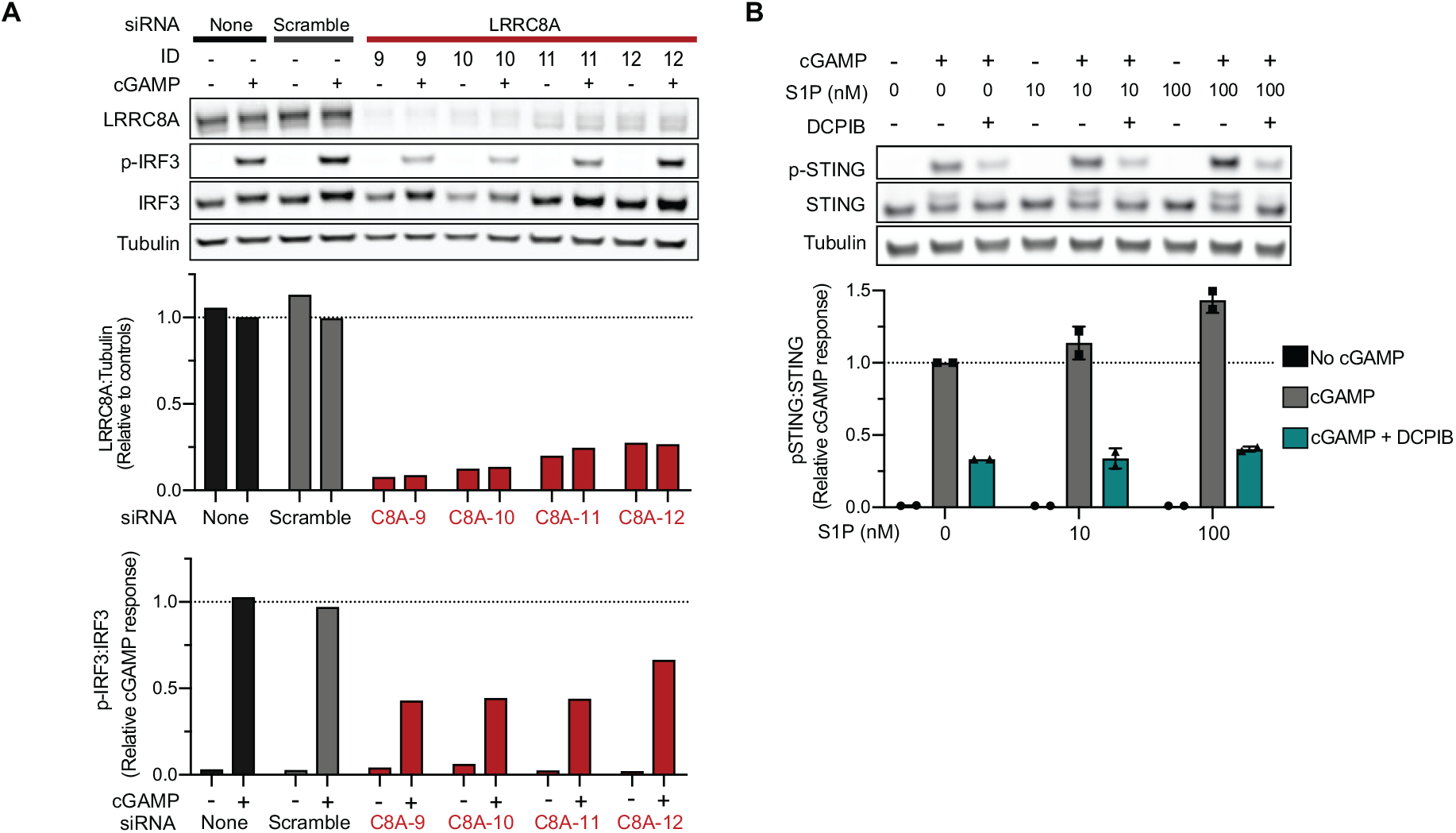
LRRC8A-Containing Channels Are the Dominant cGAMP Importer in Primary Human Endothelial Cells. (A) Effect of LRRC8A on extracellular cGAMP signaling in primary HUVEC cells. Following siRNA knockdown of LRRC8A expression in HUVEC (4 d), cells were treated with cGAMP (100 μM) for 2 h. Western blots shown with quantification (n = 1). (B) Effect of S1P and DCPIB on extracellular cGAMP signaling in HUVEC cells. Cells were treated with cGAMP (50 μM), with or without S1P (10 or 100 nM) and DCPIB (20 μM) for 1 h. Representative Western blots shown with quantification of mean ± SD (n = 2 biological replicates).

Given the modulation of LRRC8A-mediated cGAMP import that we achieved using S1P and DCPIB in the microvasculature line, we tested whether these same tools could effectively regulate cGAMP import in primary human endothelial cells. Consistent with our earlier findings, extracellular cGAMP signaling was increased in HUVEC cells, here by up to 1.4-fold, in the presence of physiological S1P concentrations (Figure 7B). DCPIB again inhibited both basal and S1P-potentiated cGAMP import. In total, these data establish pharmacologic modulation of the LRRC8A channel as a potential strategy to control cGAMP import into human vasculature.

## DISCUSSION

Here we report identification of the LRRC8A:C heteromeric channel as the second known transporter of extracellular cGAMP. The LRRC8 family was recently discovered to form the channels responsible for VRAC activity after three decades of active searching (Cahalan and Lewis, 1988; Grinstein et al., 1982; Qiu et al., 2014; Voss et al., 2014). With the molecular identity of this channel now known, detailed mechanistic studies are now possible. Beyond small anions, LRRC8A channels transport metabolites, neurotransmitters, and small molecule drugs (Lee et al., 2014; Lutter et al., 2017; Planells-Cases et al., 2015). It is becoming clear that VRAC’s namesake function in regulation of cell volume represents only a subset of the broad physiological roles the channels play. LRRC8A has been linked to cellular development, migration, and apoptosis, as well as cell-to-cell communication by regulating insulin release and mediating excitatory astrocyte-neuron signaling (Chen et al., 2019; Osei-Owusu et al., 2018). We identify immunostimulatory 2’3’-cyclic dinucleotides as a new class of substrates transported by LRRC8A-containing channels.

Structural studies have now revealed notable similarities and differences between two complexes that transport cGAMP: LRRC8A-containing channels and connexins. In the solved homomeric structures of LRRC8A and connexin Cx26, both form hexameric complexes with the four alpha helices of the respective transmembrane regions forming highly analogous secondary structures (Deneka et al., 2018; Maeda et al., 2009). This homology supports the idea that substrates of these complexes, such as cGAMP, may overlap. In the case of gap junctions, two connexin complexes on neighboring cells associate to form a channel between the two plasma membranes that allows direct cytosol-to-cytosol transport. In contrast, a single LRRC8A complex forms a channel that links the cytosol to the extracellular space and is gated by a narrow extracellular pore. While gap junction cGAMP transfer enables rapid signal transduction in tightly packed cell matrices (Ablasser et al., 2013b), LRRC8A-mediated transport is poised to enable longer-range cGAMP communication through the extracellular milieu to non-adjacent or migrating cells. In this manner, these large channel complexes likely play complementary and non-overlapping roles in paracrine cGAMP signaling.

Once the LRRC8A channel is open, cGAMP travels down its chemical gradient across the plasma membrane. Indeed, other than serving as a cGAMP importer, we demonstrated that the LRRC8A channel, when activated, exports cGAMP out of HEK cells that overexpress cGAS. Therefore, LRRC8A is first reported cGAMP exporter. The role that the LRRC8A channel plays as an exporter in cell types with high endogenous cGAMP warrants further investigation.

The identification of a new substrate class for LRRC8A channels raises the question of how one family of channels can have such versatile roles. While LRRC8A-mediated VRAC function is reported to be ubiquitous, and we observed a contribution of LRRC8A to cGAMP signaling in all cell lines evaluated, it is evident that the exact subunit composition of the heteromeric complex dictates substrate specificity and physiologic role. We demonstrate that LRRC8C, when complexed with the obligatory LRRC8A subunit, directly transports cGAMP. While LRRC8C and LRRC8E exhibit high homology (Abascal and Zardoya, 2012), expression of LRRC8E is restricted. We observed little to no role for LRRC8E in cGAMP transport; however, it is conceivable that LRRC8A:E complexes could contribute to CDN transport when sufficiently abundant. The presence of LRRC8D inhibits cGAMP transport, possibly because it forms non-productive complexes with LRRC8A or LRRC8A:C. Given the requirement of a specific LRRC8A heteromeric complex and differential expression profiles of the paralogs across cell types, we noticed that human vasculature cells express high levels of *LRRC8A* and *LRRC8C*, but low levels of *LRRC8D*. Indeed, an immortalized microvasculature cell line and primary human endothelial cells use LRRC8A channels as a dominant cGAMP import mechanism. We predict that other cell lineages that share this LRRC8 expression profile would also transport cGAMP via this route.

LRRC8A channels represent the first identification of a dominant cGAMP transport mechanism in primary human cells, since the previously identified cGAMP importer SLC19A1 plays limited roles in tested primary cells (Ritchie et al., 2019b, 2019a). Our identification of LRRC8A channels as the dominant cGAMP importer in vasculature cells also provides a molecular mechanism for sensing other 2’3’-CDNs. Vascularization of tumors is required to meet high metabolic demand and allows for tumor outgrowth, and therefore has been a target for anti-cancer therapeutic development for decades. Tumor vasculature-disrupting agents have shown promising results in treatment of solid tumors by shutting down blood flow and causing massive tumor necrosis (Tozer et al., 2005). It was recently shown that the ability of the cGAMP analog 2’3’-CDA^S^ to induce tumor necrosis depends on activation of STING in stromal cells and, moreover, that signaling between the stromal and hematopoietic compartments is beneficial for generation of anti-cancer immunity (Francica et al., 2018). With 2’3’-CDA^S^ currently undergoing evaluation in clinical trials for treatment of metastatic cancers, establishing how the investigational new drug reaches effector cells will aid understanding of its efficacy and potential toxicity. Our results suggest that LRRC8A channels likely represent a major uptake mechanism of 2’3’-CDA^S^ injected into tumor vessels or accessible to the microvasculature. LRRC8A knockout mice are poorly viable (Kumar et al., 2014), therefore, future studies should focus on development of specific drug-like molecules to regulate channel activity *in vivo.* Building upon the discoveries described here, modulation of LRRC8A:C channel activity represents a novel pharmacologic strategy to selectively increase or decrease extracellular CDN to STING signaling, especially in vasculature, and may hold the potential to tune downstream innate immune activation and elicitation of anti-tumor responses.

## ACKNOWLEDGMENTS

We thank all Li Lab members for their insightful comments and discussion throughout the course of this study. Flow cytometry analysis for this project was done on instruments in the Stanford Shared FACS Facility. Data was collected on an instrument in the Shared FACS Facility obtained using NIH S10 Shared Instrument Grant S10RR027431-01. L.J.L. thanks the Stanford Graduate Fellowship, ARCS Foundation, and Kimball Foundation for support. This research was supported by the National Institutes of Health grants DP2CA228044 (L.L.), DP2HD084069 (M.C.B.), and DOD grant W81XWH-18-1-0041 (L.L.).

## AUTHOR CONTRIBUTIONS

All authors designed the studies and discussed the findings in the manuscript. L.J.L., X.W., R.E.M., V.B., and G.T.H. performed experiments and analyzed data. C.R. and J.A.C. provided technical support. L.J.L. and L.L. wrote the manuscript.

## DECLARATION OF INTERESTS

The authors declare no competing interests.

## EXPERIMENTAL METHODS

### Cell culture

Unless otherwise noted, cell lines were obtained from ATCC. U937 cells were maintained in RPMI (Cellgro) supplemented with 10% heat-inactivated FBS (Atlanta Biologicals) and 1% penicillin-streptomycin (Gibco). HEK293 cells were maintained in DMEM (Cellgro) supplemented with 10% FBS and 1% penicillin-streptomycin.

Telomerase-immortalized human microvascular endothelium (TIME) cells line were maintained in vascular cell basal media supplemented with microvasculature endothelial cell growth kit-VEGF (ATCC) and 0.1% penicillin-streptomycin. Human umbilical vein endothelial cells (HUVEC) pooled from multiple donors were purchased from Lonza and maintained in EGM-2 supplemented growth media (Lonza). All cells were maintained in a 5% CO_2_ incubator at 37 °C.

### Reagents and antibodies

2’3’-cyclic-GMP-AMP (cGAMP) was synthesized and purified in-house as previously described (Ritchie et al., 2019a). DMXAA, 2’3’-bisphosphorothioate-cyclic-GMP-AMP (2’3’-cG^S^A^S^MP), 2’3’-cyclic-di-AMP (2’3’-CDA), 2’3’-bisphosphorothioate-cyclic-di-AMP (2’3’-CDA^S^), 3’3’-cyclic-GMP-AMP (3’3’-cGAMP), 3’3’-cyclic-di-AMP (3’3’-CDA), and 3’3’-cyclic-di-GMP (3’3’-CDG) were purchased from Invivogen and reconstituted in endotoxin-free water. Sphingosine-1-phosphate (d18:1) was purchased from Cayman Chemical. 4-[(2-Butyl-6,7-dichloro-2-cyclopentyl-2,3-dihydro-1-oxo-1*H*-inden-5-yl)oxy] butanoic acid (DCPIB) was purchased from Tocris Bioscience and reconstituted to 50 mM in DMSO. Antibodies and dilutions used for Western blotting are listed in Table S1.

### Generation of CRISPR edited cell lines

LentiCRISPR v2 (Addgene) was used as the 3^rd^-generation lentiviral backbone for all knockout lines. Sequences for all sgRNAs used in this study are listed in Table S2. The guide sequences were cloned into the lentiviral backbone using the Lentiviral CRISPR Toolbox protocol from the Zhang Lab at MIT (Sanjana et al., 2014; Shalem et al., 2014). Lentiviral packaging plasmids (pHDM-G, pHDM-Hgmp2, pHDM-tat1b, and RC/CMV-rev1b) were purchased from Harvard Medical School. 500 ng of the lentiviral backbone plasmid containing the guide sequence and 500 ng of each of the packaging plasmids were transfected into HEK 293T cells using FuGENE 6 transfection reagent (Promega). The viral media was exchanged after 24 hours, harvested after 48 hours and passed through a 0.45 μm filter. U937 cells were transduced by spin infection in which cells were suspended in viral media with 8 μg/mL polybrene (Sigma Aldrich), centrifuged at 1000 x g for 1 hour, then resuspended in fresh media. HEK293 and TIME cell lines were reverse transduced by trypsinizing adherent cells and adding cell suspensions to viral media supplemented with 8 μg/mL polybrene. All lines were put under relevant antibiotic selection beginning 72-hours post-transduction and lasting until control (untransduced) cells completely died.

### Cell viability measurement

U937 cells were treated with the indicated concentrations of cGAMP for 24 or 48 hours in duplicate. Cell density and viability was determined by hemocytometer and trypan blue staining.

### U937 *SLC19A1*^*−/−*^ CRISPR knockout library generation

Design of the whole-genome CRISPR sgRNA library was previously described by Morgens et al. (2017). Briefly, a whole-genome library of exon-targeting sgRNAs were designed, with the goal of minimizing off-target effects and maximizing gene disruption. The top 10 sgRNA sequences for each gene were included in the library, along with thousands of safe-targeting and non-targeting negative controls. The library was cloned into a lentiviral vector, pMCB320, which also expresses mCherry and a puromycin resistance cassette. The U937 Cas9-*SLC19A1*^*−/−*^ line was generated by lentiviral transduction as described above. Knockout of *SLC19A1* was confirmed by genomic DNA sequencing (Figure S1A) and Cas9 function was confirmed by the efficiency of knocking out GFP expression using a sgRNA sequence against GFP confirmed by using a sgRNA sequence against GFP (Figure S1B). U937 Cas9-*SLC19A1*^*−/−*^ cells were infected with the lentiviral CRISPR sgRNA library and selected with puromycin.

### CRISPR screen

The U937 CRISPR knockout library line was grown in 4 spinner flasks (1 L), with 2 flasks serving as untreated controls and 2 flasks receiving cGAMP treatment.

Throughout the screen all of the samples were split daily to keep the cell density at 250 million cells per 500 mL, which corresponded to 1,000 cells per guide in the untreated samples. The experimental samples were treated daily with enough cGAMP to reduce treated cell fold growth by 50% as compared to the control samples. Initial treatments began at 15 μM for two days and were then titrated to 30 μM. After a difference of ten population doublings was achieved between experimental and control samples (12 days), the genomic DNA was extracted using a Qiagen Blood Maxi Kit. The library was sequenced using a NextSeq 500/550 Mid Output v2 kit (Illumina). The experimental and control samples were compared using casTLE (Morgens et al., 2016), available at https://bitbucket.org/dmorgens/castle. The algorithm determines the likely effect size for each gene, as well as the statistical significance of this effect.

### Sequencing of gDNA to confirm CRISPR editing

gDNA was isolated from cell lines using the QIAamp DNA Mini Kit (Qiagen). Sequences of each gene loci flanking the sgRNA target sequence were PCR amplified. PCR products were purified and submitted for Sanger sequencing analysis. Control (unedited) and experimental (edited) sequence traces were analyzed using the Inference of CRISPR Edits (ICE) version2 software tool (Synthego). ICE knockout scores represent the proportion of sequences in which a coding frameshift or >21-bp insertion/deletion are detected.

### Electroporation of STING agonists

U937 cells were resuspended in 100 μL electroporation solution (90 mM Na_2_HPO_4_, 90 mM NaH_2_PO_4_, 5 mM KCl, 10 mM MgCl_2_, 10 mM sodium succinate) with the appropriate concentration of STING agonist. Cells were then transferred to a cuvette with a 0.2 cm electrode gap (Bio-Rad) and electroporated using program U-013 on a Nucleofector II device (Lonza). Following electroporation, cells were transferred to media and cultured as indicated. Cell lysates were then harvested for analysis.

### Co-expression of mSTING and agonist treatment with varied osmolarity

HEK293 cell lines were seeded in 12-well plates at 100,000 total cells in 2 mL media one day before transfection. Each well was transfected with 3 uL FuGENE 6 reagent (Promega) and 130 ng pcDNA3-mouseSTING in line with manufacturer’s protocols.

Isotonic (130 mM NaCl, 6 mM KCl, 1 mM MgCl_2_, 1.5 mM CaCl_2_, 10 mM glucose, 10 mM HEPES, pH 7.4, ~300 mOsm) or hypotonic buffer (105 mM NaCl, 6 mM KCl, 1 mM MgCl_2_, 1.5 mM CaCl_2_, 10 mM glucose, 10 mM HEPES, pH 7.4, ~240 mOsm) containing either DMXAA (250 μM) or cGAMP (100 μM) was used to treat cells for 1 h.

### Stimulation of cGAMP synthesis and measurement of export in HEK293 cells

HEK293 cell lines were seeded in PurCol-coated (Advanced BioMatrix) 6-well plates at 300,000 total cells in 2 mL media one day before transfection. At the start of the experiment, media was gently removed and replaced with complete DMEM supplemented with ENPP1 inhibitor STF-1084 for 50 μM final. Cells were then transfected with 1500 ng pcDNA-FLAG-HA-sscGAS plasmid complexed with FuGene 6 reagent (Promega) according to manufacturer’s instructions or treated with FuGene 6 alone as a negative control. After 24 hours of incubation, media from each condition was harvested and cells were immediately incubated for 20 minutes with hypotonic buffer (60 mM NaCl, 6 mM KCl, 1 mM MgCl_2_, 1.5 mM CaCl_2_, 10 mM glucose, 10 mM HEPES, pH 7.4, 155 mOsm). Following collection of buffer, media and buffer samples were centrifuged at 1000 x g for 10 minutes and supernatants collected. Extraction of cGAMP was then performed using HyperSep Aminopropyl SPE columns (ThermoFisher Scientific) and sample submitted for mass spectrometry quantification of cGAMP as previously described (Carozza et al., 2020).

### Electrophysiology

Endogenous VRAC currents from HEK293 cells were measured by whole-cell patch-clamp. Cells were transferred from culture dishes to the electrophysiology recording chamber following treatment with 0.5 mg/mL trypsin (Sigma-Aldrich) for 2 minutes.

External recording solution contained 88 mM NaCl, 10 mM HEPES with either 110 mM mannitol (isotonic, 300 mOsm/kg) or 30 mM mannitol (hypotonic, 230mOsm/kg), pH 7.4 by NaOH. The internal (pipette) solution contained 130 mM CsCl, 10 mM HEPES, 4 mM Mg-ATP, pH 7.3. Borosilicate glass micropipettes (Sutter Instruments, Novato, CA) were pulled and fire-polished to a tip diameter with a typical resistance of 1.5–3.0 MΩ. Data were acquired using an Axopatch-200B amplifier (Axon Instruments, Union City, CA) and InstruTECH ITC-16 interface (HEKA Instruments, Holliston, MA), with a sampling rate of 5 kHz and filtering at 1 kHz. Igor Pro (WaveMetrics, Portland, OR) software was used for stimulation and data collection. Currents were measured in response to voltage ramps from −150 to 150 mV over 1.0 s, with an inter-ramp interval of 10 s and a holding potential of 0 mV. All recordings were carried out at room temperature (20 – 22 ºC).

### Lipofection of cGAMP

Lipofectamine 3000 (Thermo Fisher Scientific) was diluted into Opti-MEM (3 μL into 50 μL), P3000 was diluted into Opti-MEM (2 μL into 50 uL) in the absence or presence of cGAMP (for 1 μM final in experiment), and then dilutions were combined and incubated for 15 minutes. Lipid complexes (100 μL) were added dropwise to plated cells in 900 uL media. In parallel, free cGAMP was added for 1 μM final in the absence of lipid reagent.

### Inducible expression of LRRC8A and LRRC8C

Gene fragments encoding LRRC8A (Uniprot Q8IWT6) and LRRC8C (Uniprot Q8TDW0) were commercially synthesized and cloned into plasmids (Twist Bioscience). Synonymous DNA mutations were introduced by QuickChange to ablate downstream Cas9-sgRNA targeting at each site (LRRC8A: GGATCCTGAAGCCGTGGT to G<u>C</u>AT<u>TT</u>T<u>A</u>AA<u>A</u>CC<u>A</u>TGGT, LRRC8C: GTTATGAGCGAGCCCTCCAC to G<u>C</u>TA<u>C</u>GA<u>A</u>CG<u>C</u>GC<u>GT</u>T<u>A</u>CA<u>T</u>). Additionally, QuickChange was used to generate a sequence with a LRRC8A-R103L encoding substitution (CGG to CTG). A parent pLVX-TetOne plasmid (Takara Bio) was modified by 1) insertion of a GGSG-FLAG encoding sequence to flank the multiple cloning site and 2) insertion of SV40 promoter and hygromycin resistance factor encoding sequences were cloned in following the TetOn 3G element, yielding a pLVX-TetOne-FLAG-Hygro plasmid. LRRC8A, LRRC8A-R103L, and LRRC8C DNA fragments (all with ablated sgRNA sites) were then cloned into pLVX-TetOne-FLAG-Hygro backbone by Gibson assembly. Lentiviral packaging of constructs was performed as described above and reverse transduced into LRRC8A^−/−^ or LRRC8C^−/−^ TIME cells. Cells were selected with hygromycin for 2 weeks. Doxycycline was added to cultures for 36 hours at the indicated concentrations before each experiment to induce expression.

## S1P RECONSTITUTION

Lipid was solubilized in warm methanol, aliquoted, then solvent evaporated under nitrogen before storage at −20*C. At the start of each experiment, S1P was freshly reconstituted to 100 μM with 4 mg/mL fatty acid-free human serum albumin carrier protein (Millipore Sigma) in PBS.

### siRNA knockdown of *LRRC8A* in HUVEC

Four ON-TARGETplus siRNAs for knockdown of *LRRC8A* were purchased from Dharmacon, along with non-targeting control siRNA. HUVEC cells were seeded the night before transfection in 6-well plates at 2.5×10^5^ total cells in 2 mL media, with a change to fresh media on the day of knockdown. Following manufacturer’s instructions DharmaFECT4 (6 uL) was complexed with siRNA (10 uL of 5 uM stock) in Opti-MEM to yield 200 uL of complexes then used to transfect cells. Cells were split on day three post-transfection and tested for cGAMP response on day four.

### Statistical analysis

All statistical analyses were performed using GraphPad Prism 8.3.1. For all experiments involving Western blots, densitometric measurements of protein bands were made using ImageJ 1.52a. For comparison of values attained within one blot (Figure 2A), significance was calculated by two-tailed *t*-test, assuming a Gaussian distribution. In cases where biological replicates of independent experiments represent paired sets of blots (Figures 3-5), significance was calculated by ratio paired *t*-test assuming a Gaussian distribution.

---

**Figure S1.**
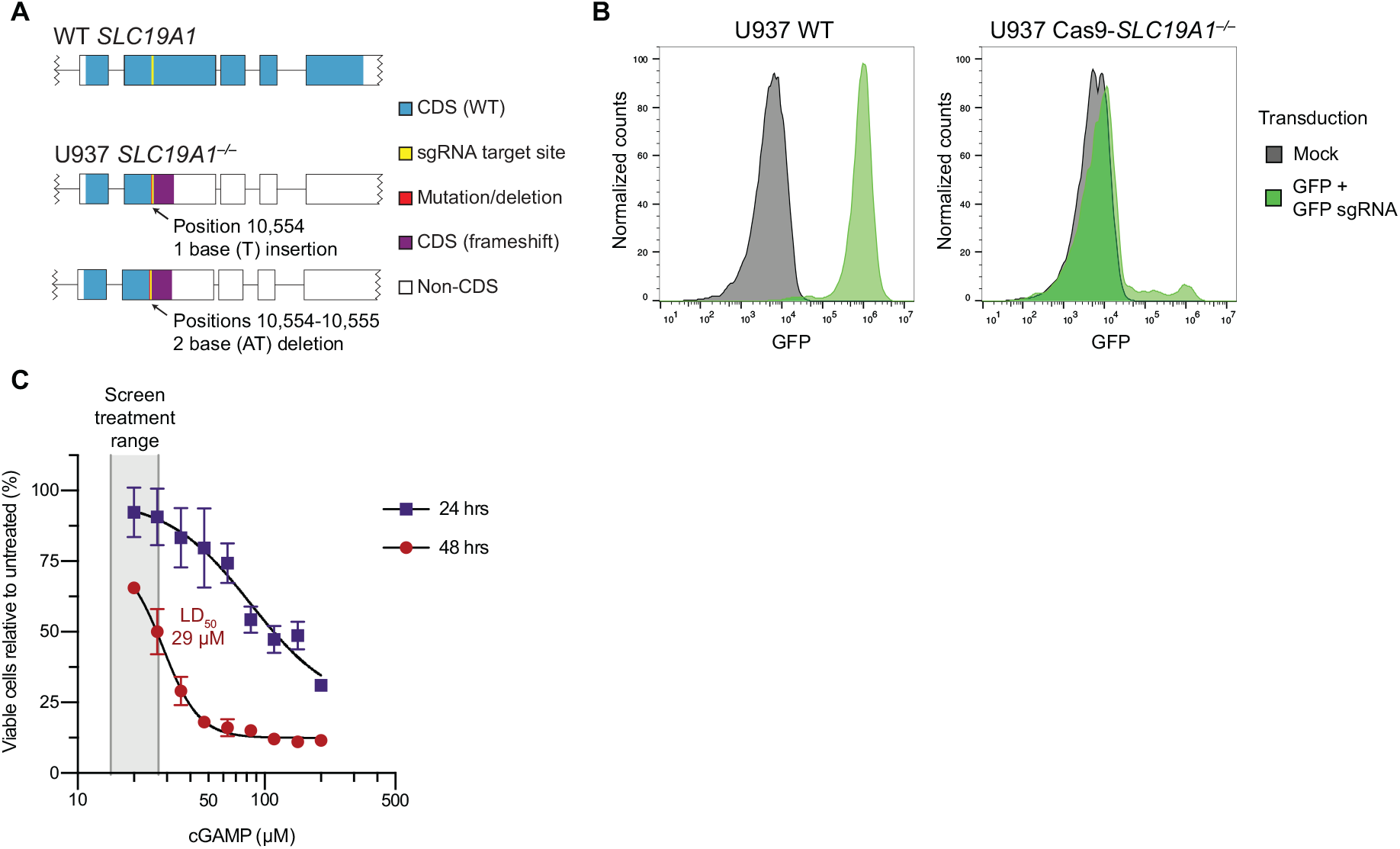
A Genome-Wide CRISPR Screen Identifies LRRC8A as a Positive Regulator of Extracellular cGAMP-Mediated STING Pathway Activation, Related to Figure 1. (A) Diagram of mutations in *SLC19A1* loci with predicted coding effects in the U937 Cas9-*SLC19A1*^*−/−*^ line. Genomic DNA was isolated from cells and DNA flanking the *SLC19A1* sgRNA target site was sequenced. Editing is present in both alleles beginning at position 10,554 of the *SLC19A1* gene, with allele #1 containing an insertion and allele #2 containing a deletion. Both mutations result in a coding frameshift in exons 2 and 3, terminating in a premature stop in exon 3. Boxes represent coding exons of *SLC19A1* and horizontal lines represent introns. (B) Plot of GFP expression measured as a test of Cas9 editing efficiency in the U937 Cas9-*SLC19A1*^*−/−*^ line. U937 WT or U937 Cas9-*SLC19A1*^*−/−*^ cells were untreated or transduced with lentivirus encoding GFP expression and GFP-targeting sgRNA. Transduced cells express GFP in the absence of Cas9, while Cas9 expression results in CRISPR knockout of GFP. Cells were analyzed for GFP expression by flow cytometry and data are displayed as histograms of ~10,000 events from each condition. (C) Plot of percent live U937 *SLC19A1*^*−/−*^ cells in response to cGAMP treatment after 24 or 48 h. Cells were counted by hemocytometry. Data are mean ± SD (n = 3 biological replicates at 24 h, n = 2 biological replicates at 48 h).

**Figure S2.**
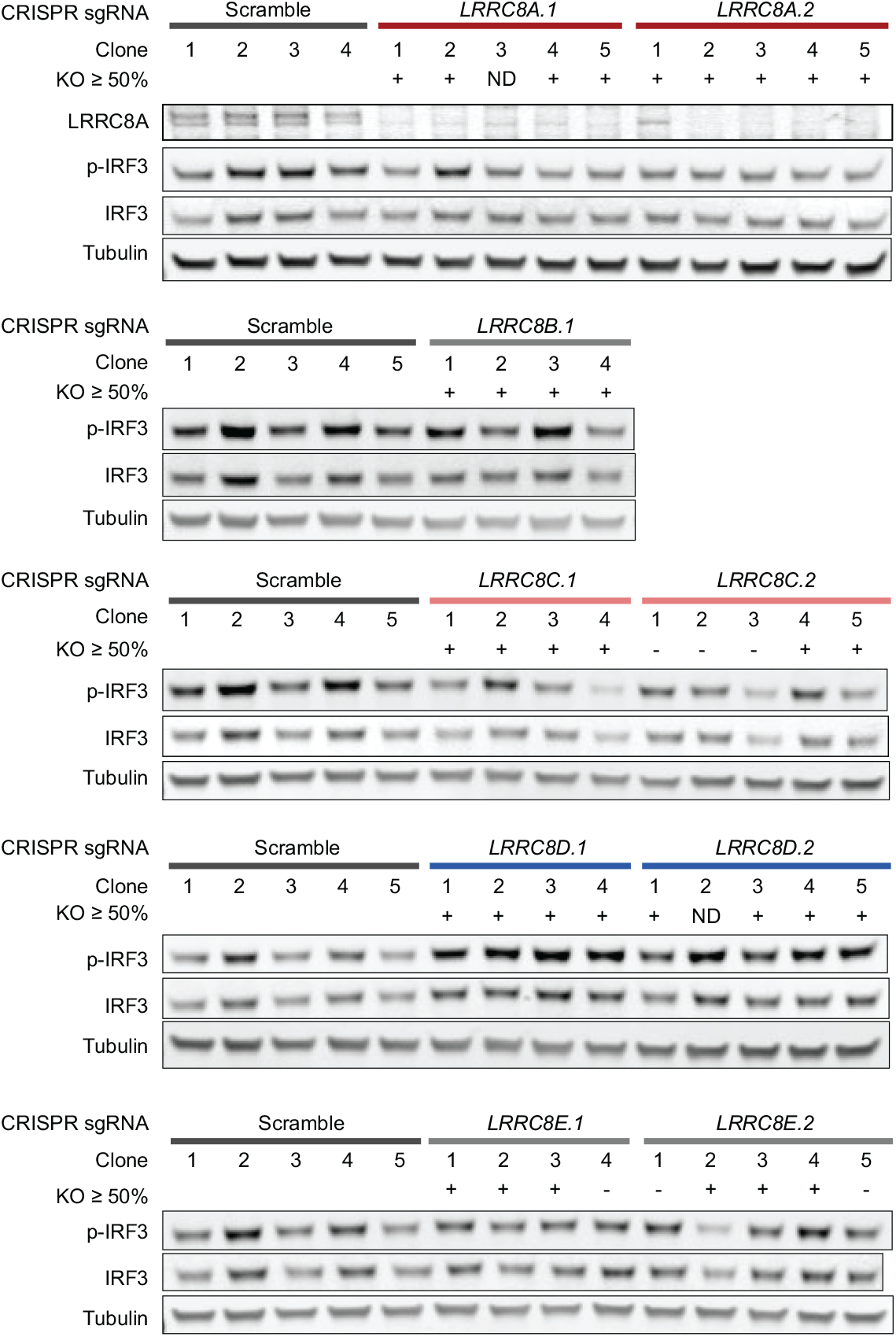
LRRC8A and LRRC8 Paralogs Differentially Facilitate cGAMP Import, Related to Figure 2. U937 *SLC19A1*^*−/−*^ clones expressing scramble or *LRRC8A-E* sgRNAs were treated with 100 μM cGAMP for 4 h and signaling was measured by Western blot. Validation of CRISPR editing was confirmed by gDNA sequencing and Inference of CRISPR Edits (ICE) analysis. ICE knockout (KO) scores represent the proportion of sequences with a frameshift or >21 base pair insertion/deletion, with scores ≥ 50 found upon heterozygous or homozygous gene knockout. Not determined (ND) represents poor sequence traces preventing analysis. The same scramble clone lysates were run with each *LRRC8* knockout group, enabling the relative normalization of confirmed (KO ≥ 50) clones displayed in Figure 2A.

**Figure S3.**
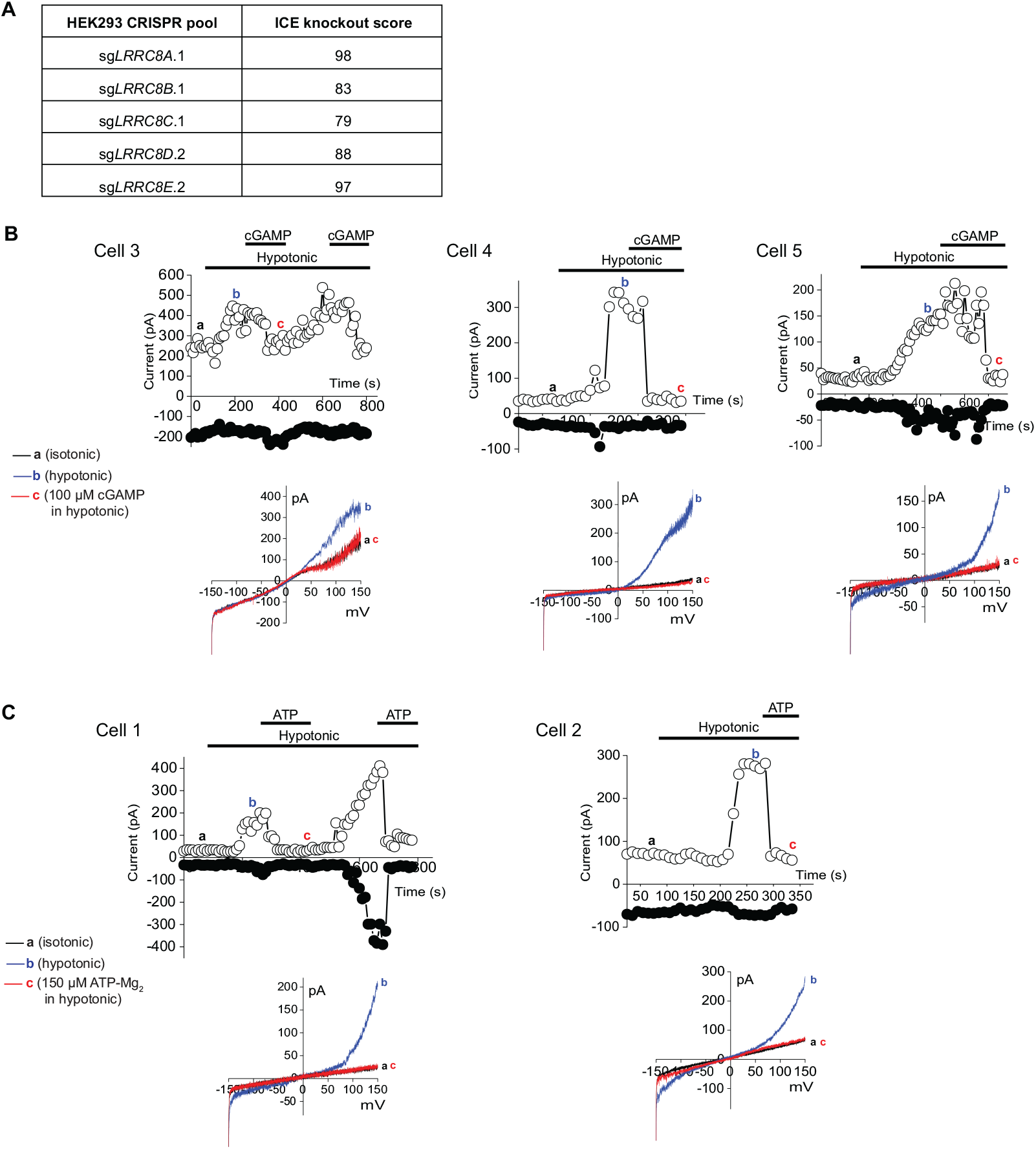
LRRC8A-Containing VRAC Channels Directly Transport cGAMP, Related to Figure 3. (A) Table reporting ICE knockout scores for HEK293 knockout pools, as determined from gDNA sequencing at the corresponding locus. Knockout scores represent the proportion of sequences with a frameshift or >21 base pair insertion/deletion. (B) Representative whole-cell patch clamp current traces measured in HEK293 cells upon hypotonic stimulation in the absence or presence of cGAMP (100 μM). Overlays of the current-voltage relationship at the indicated points are also displayed. Traces for cells 1 and 2 are displayed in Figure 3C. (C) Representative whole-cell patch clamp current traces measured in HEK293 cells upon hypotonic stimulation in the absence or presence of ATP (150 μM). Overlays of the current-voltage relationship at the indicated points are also displayed.

**Figure S4.**
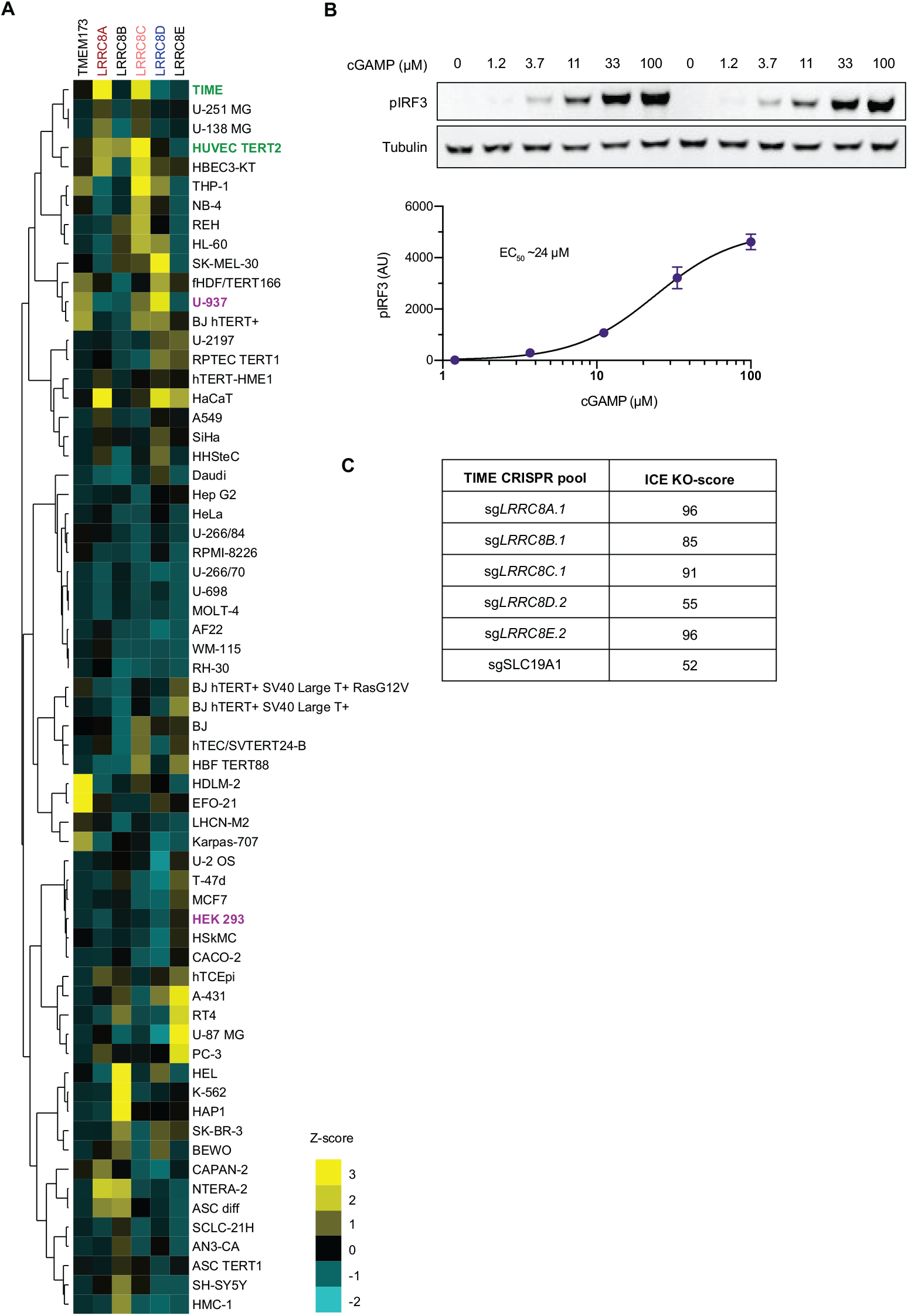
The LRRC8A:C Channel Is the Dominant cGAMP Importer in Microvasculature Cells, Related to Figure 4. (A) Heatmap of RNA transcript levels of *TMEM173* (STING) and *LRRC8* paralogs across common cell lines. Publicly available RNAseq data were downloaded from The Human Protein Atlas (proteinatlas.org). Minimum detectable expression was set to NX = 1 and Z-scores were then calculated within each transcript (observed _transcript x_ – average _transcript x_) / standard deviation transcript x. Hierarchical clustering was applied for display of cells based on similarity of transcript profiles. Vasculature lineages are annotated in green text, and other lines studied in the paper are annotated in purple. (B) Dose-response determination of extracellular cGAMP in TIME cells. Wild-type TIME cells were treated with cGAMP at the indicated concentrations for 2 h and signaling measured by Western blot. Data are mean ± SD (n = 2 technical replicates). (C) Table reporting ICE knockout scores for TIME knockout pools, as determined from gDNA sequencing at the corresponding locus. The knockout score represents the proportion of sequences with a frameshift or >21 base pair insertion/deletion.

**Figure S5.**
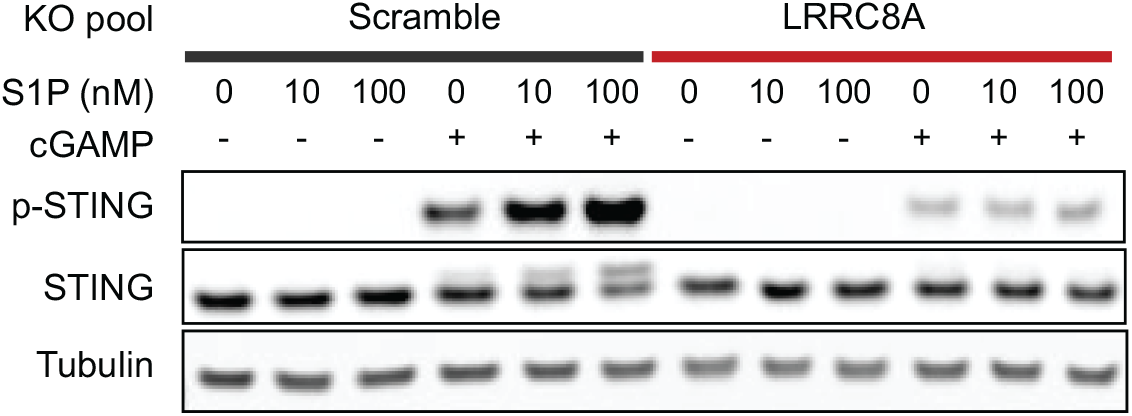
cGAMP Import by the LRRC8A:C Channel Can Be Potentiated by Sphingosine-1-Phosphate and Inhibited by DCPIB, Related to Figure 5. (A) TIME cells were treated with sphingosine-1-phosphate (S1P) with or without cGAMP for 1 h and STING pathway activation was assessed by Western blot (n = 1).

**Table S1.**
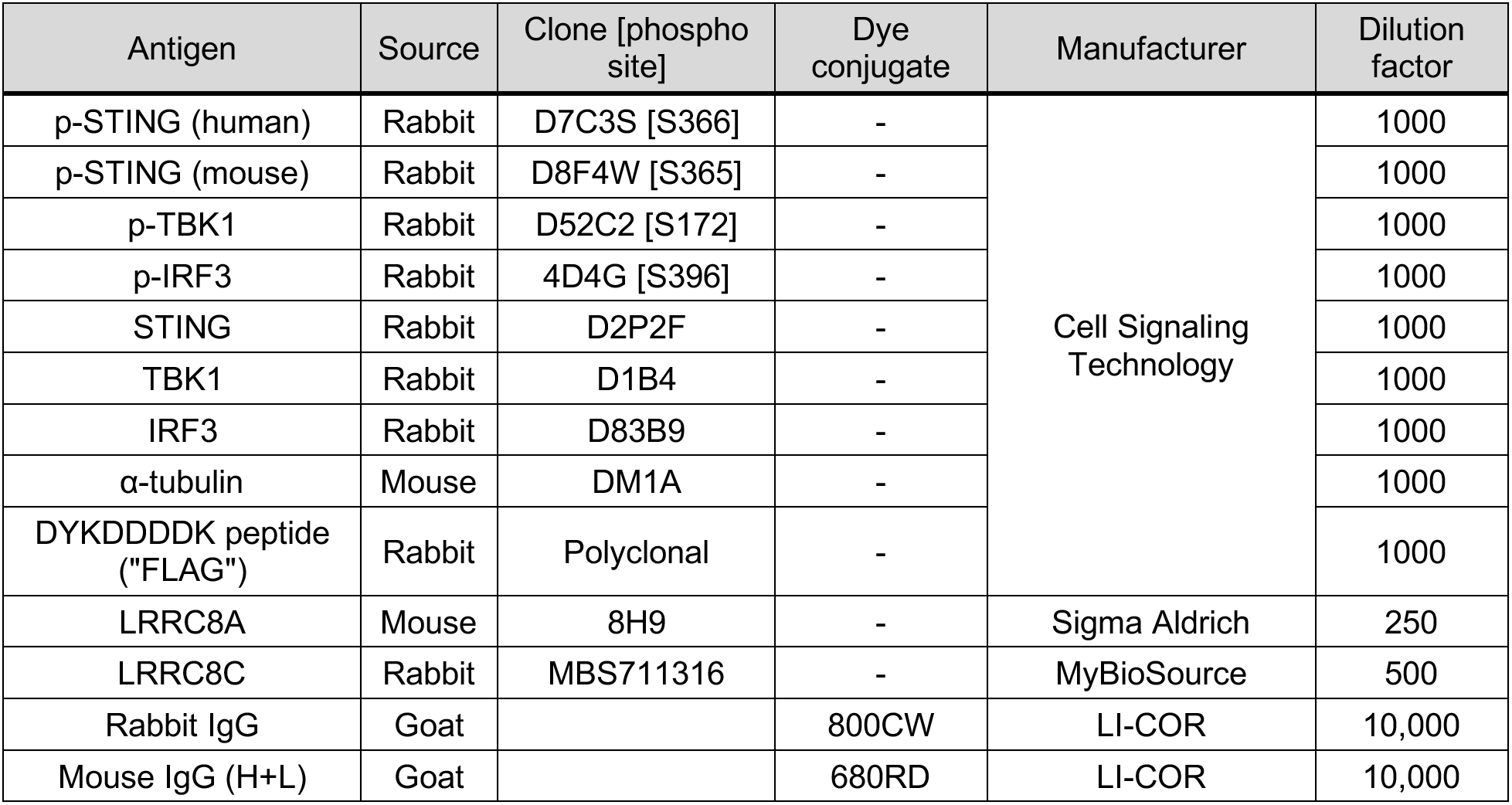
Antibodies Used in This Study, Related to Experimental Methods.

**Table S2.**
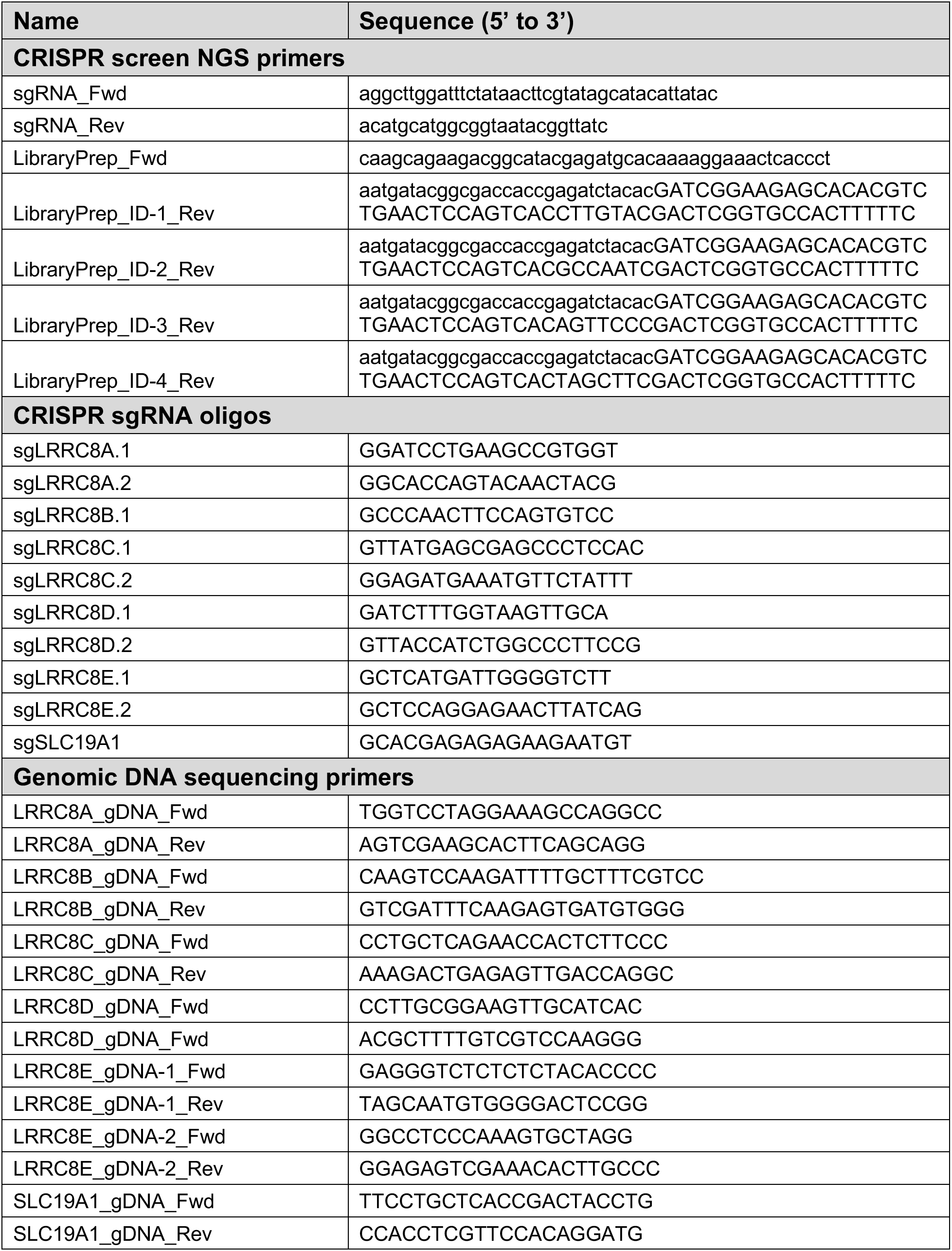
Oligonucleotides Used in This Study, Related to Experimental Methods.

